# PAMD-Ch17, a Polymeric Analog of Plerixafor, Induces Mitochondrial Dysfunction in T-ALL Cells Independent of CXCR4

**DOI:** 10.1101/2025.05.28.656643

**Authors:** Calvin Lam, Arjun Dhir, Chinmay M. Jogdeo, Sipra Panda, Ekta Kapoor, Siyuan Tang, Victor Rivero, Erin M. McIntyre, Donald W. Coulter, Peng Xiao, Adrian R. Black, Kyle Hewitt, Samantha A Swenson, Svetlana Romanova, David Oupicky, R. Katherine Hyde

**Author notes:** Corresponding Author R. Katherine Hyde, PhD, 985879 Nebraska Medical Center, Omaha NE 68198.

## Abstract

PAMD-Ch17 is a polymer composed of the CXCR4 inhibitor AMD3100/Plerixafor with a cholesterol modification. In previous work, we showed that PAMD-Ch17, but not AMD3100, induces cell death and differentiation in mouse Acute Myeloid Leukemia cells. To investigate the mechanism of PAMD-Ch17’s novel anti-leukemic effects, we tested PAMD-Ch17 against a panel of human leukemia cell lines and found that PAMD-Ch17 is effective against a variety of acute leukemias, with T-ALL cell lines being highly sensitive. Surprisingly, *CXCR4* knock out T-ALL cells were equally sensitive to PAMD-Ch17. Using a fluorescently tagged PAMD-Ch17, we found that the drug colocalized to the mitochondria. We also found that PAMD-Ch17 induced changes in expression of genes related to mitochondrial function, increased levels of mitochondrial superoxide, and decreased mitochondrial membrane potential. Using seahorse assays, we found that PAMD-Ch17 decreased baseline oxygen consumption, ATP production, and proton leakage. PAMD-Ch17 also decreased baseline extracellular acidification rate, indicating a decrease in overall metabolism. In mouse primary T-ALL but not healthy bone marrow cells, PAMD-Ch17 induced both mitochondrial superoxide and cell death. Using human bone marrow organoids, we found that PAMD-Ch17 induced mitochondrial superoxide and cell death in human primary T-ALL cells, but not in healthy stromal and hematopoietic cells. Collectively, our results indicate that PAMD-Ch17 has anti-leukemic effects against T-ALL cells but not healthy cells, likely mediated through a CXCR4 independent, mitochondrial based mechanism. These findings support further development of PAMDs as potential therapeutics for patients with T-ALL.

**KEY POINTS:**

- PAMD-Ch17, a polymeric drug based on AMD3100/Plerixafor, has novel anti-leukemic activities against T-ALL that are independent of CXCR4 inhibition.
- PAMD-Ch17 induces increased mitochondrial superoxide and cell death in primary ALL cells, but not healthy bone marrow cells.

## INTRODUCTION

T-cell acute lymphoblastic leukemia (T-ALL) is a blood cancer that is characterized by rapid, aggressive expansion of immature T cells. These immature cells outcompete the native bone marrow and hematopoietic cells, leading to anemias, lymphadenopathy, bruising, hepatomegaly, and/or splenomegaly ^1-4^. Traditional chemotherapeutics such as cytarabine and vincristine form the core of current regimens but are poorly tolerated and are associated with significant side effects ^1,2,4,5^. Therapeutics that target specific mutations are sparse with utility limited to only a subset of patients. There is a strong need for additional therapeutics against T-ALL.

We recently reported on a class of polymeric drugs, called PAMDs, that have potential for the treatment of leukemia. PAMDs are synthesized by Michael addition of the CXCR4 inhibitor AMD3100 and can be modified with additional functional groups such as cholesterol moieties ^6-14^. Previous work showed that PAMDs retain the ability to inhibit CXCR4 and can form nanoparticles with nucleic acids such as siRNAs and miRNAs ^6-14^. When we evaluated a PAMD nanoparticle with a cholesterol modification, PAMD-Ch17, as a vehicle for nucleic acid delivery to leukemia cells, we also found that it induces cell death and differentiation in mouse acute myeloid leukemia (AML) cells. Interestingly, neither AMD3100 nor other structurally unrelated CXCR4 inhibitors have similar anti-leukemic effects, consistent with previous reports ^15-17^. These findings imply that PAMD-Ch17 has additional activities unrelated to the inhibition of CXCR4.

Here we show that PAMD-Ch17 has anti-leukemic effects against T-ALL cell lines. In addition, we found that while PAMD-Ch17 effectively inhibits CXCR4 in Jurkat cells, it does not require the receptor for its anti-leukemic effects. Rather, we found that PAMD-Ch17 colocalized with the mitochondria, and induced mitochondrial stress, damage and dysfunction. Importantly, we found that PAMD-Ch17 induced mitochondrial stress and death in mouse primary T-ALL cells and human primary T-ALL cells, but not in mouse or human healthy bone marrow cells. Altogether, our data demonstrate that PAMD-Ch17 induces mitochondrial dysfunction selectively in leukemia cells, providing an explanation for its novel anti-leukemic effects, and highlighting the drug’s therapeutic potential as an anti-leukemic agent.

## METHODS

### Synthesis of PAMD-Ch17

PAMD-Ch17 was synthesized and characterized as previously described ^7,10,12-14^. The molecular weight of PAMD-Ch17 was 16.7 kg/mol and the cholesterol content was 17 wt%. Fluorescently tagged PAMD-Ch17 (fPAMD) was synthesized using the fluorescent dye cy3 as previously described ^13^. Drug solutions were prepared in distilled water and kept in the dark at 4°C.

### Mice

C57BL/6 × 129SvEv F1 mice of at least 6 weeks old and of both sexes were used for experiments. Mouse primary T-ALL cells were generated by transducing bone marrow cells with the ΔEGFΔLNRΔP GFP construct provided by Dr. Warren Pear as previously described ^18,19^. Transduced bone marrow cells were transplanted into sublethalirradiated F1 mice. Mice were sacrificed at the first signs of disease, and leukemic cells harvested. Mouse primary T-ALL and healthy bone marrow cells were cultured with StemSpan SFEM I (Stemcell Technologies, Vancouver, Canada), mouse IL-3 (10ng/mL) (Grand Island, NY, Gibco), mouse IL-6 (10ng/ml) (Gibco), mouse SCF (20ng/mL) (Gibco), and 1% penicillin streptomycin glutamine (Gibco). All experiments were performed in accordance with the National Institutes of Health’s Guide for the Care and Use of Laboratory Animals. All experiments were approved by the Institutional Animal Care and Use Committee at the University of Nebraska Medical Center.

### Bone Marrow Organoid and Primary Human Leukemia Cells

Human bone marrow organoids were differentiated from human iPSCs (Gibco) as previously described ^20,21^, but with the mesodermal commitment performed in 2% O_2_, 5% CO_2_, 93% N_2_. Organoids were collected on days 10-13 and distributed into individual wells and engrafted with primary leukemia cells for 3 days prior to drug treatment ^20,21^. Human primary T-ALL cells were obtained from CureLine (San Francisco, CA) and cultured in StemSpan SFEM I (Stemcell Technologies) supplemented with 1uM UM729 (Stemcell Technologies), 10% StemSpan CD34+ Expansion Supplement (Stemcell Technologies), and 1% penicillin streptomycin glutamine (Gibco). Cells were cultured at 0.5×10^6^-10^6^ cells/ml. Prior to engraftment, leukemia cells were labeled with 1uM CellTrace Far Red (Invitrogen). 5000 viable leukemia cells were engrafted per organoid.

### Cell Lines and Culture

Molm-13, Kasumi-1, REH, Molt-4, and Jurkat cells were obtained from ATCC (Manassa, VA). SEM cells were obtained from DSMZ (Leibniz Institute, Germany). Cell lines were cultured according to supplier protocol. Briefly, Jurkat and Molt-4 cells were cultured in RPMI 1640 ATCC modification (Gibco) with 10% fetal bovine serum (FBS) supplemented with 1% penicillin streptomycin glutamine (Gibco). Cells were maintained between 0.5×10^6^-10^6^ cells/ml. Lenti-X 293T were obtained from Takara (Takara, San Jose, CA). Cells were cultured according to supplier protocol: in DMEM (Corning, Tewksbury, MA) with 10% FBS supplemented with 1% penicillin streptomycin glutamine (Gibco). Cells were maintained between 10%-80% confluency. All were cultured at 37C in a 5% CO_2_ humidified incubator.

### Whole Transcriptome Sequencing and Analysis

Jurkat cells were treated with PAMD-Ch17 (0.6uM) or control for 24 hours and harvested. RNA was isolated using TRIzol Reagent (Invitrogen, Waltham, MA). Libraries were prepared and sequenced by the University of Nebraska Medical Center (UNMC) Genomics Core Facility. Reads were trimmed via fqtrim (Phred score<30) and processed by FastQC. Differentially expressed genes (DEGs) were then identified using standard pipelines. Briefly, trimmed files were then processed using STAR as the aligner and RSEM as the annotation and quantification tool ^22,23^. Raw counts were used to obtain differentially expressed genes (DEGs) via DESeq2 ^24^.

Hg38 was used as the reference human genome. Pathway Analysis was performed using Ingenuity Pathway Analysis (Qiagen, Hilden, Germany) using a p≤0.05 to obtain pathways shown ^25^. RNA-sequencing of the *CXCR4* knockout (KO) Jurkat cells was performed and analyzed in the same manner.

### Metabolic Ability (MTT) Assay

Cells were seeded in a 96 well flat bottom plate. PAMD-Ch17/AMD3100 was prepared as 10x stock solutions in 10% dimethyl sulfoxide (DMSO) (ThermoScientific, Waltham, MA) and phosphate buffered saline (PBS) (Cytiva, Marlborough, MA). PAMD-Ch17 or AMD3100 (Selleckchem, Houston, TX) stock solution was added to each well to achieve the target dose with a final DMSO concentration of 1%, and cells were incubated for 72 hours. Prestoblue (Invitrogen) was used to assess metabolic activity, according to manufacturer’s protocol. Plates were read using Tecan Infinite M200 PRO in fluorescence top reading mode with excitation 560nm and emission 590nm. Readings were then normalized to the untreated condition.

### Microscopy

Cells were treated with the cy3 fluorescently tagged PAMD-Ch17 (fPAMD) at the indicated concentration for the indicated period of time and stained with 0.1uM Mitotracker DeepRed (Invitrogen) according to manufacturer’s recommendation, cytospun 800xg (ThermoScientific) onto charged slides (Hartfield, PA), dried, fixed in 4% paraformaldehyde (PFA) PBS, permeabilized with 0.1% Trition X-100 (Fisher BioRegeants/Thermo), and mounted with DAPI (Invitrogen ProLong Gold) before imaging on a Zeiss LSM 800 with Airyscan (Zeiss, Jena, Germany). Images were visualized using ImageJ v1.53k and colocalization quantified using Just Another Colocalization Plugin (JACoP v2.1.4) per developers recommendations ^26^. Immunofluroescence for CXCR4 was performed using anti-CXCR4 clone UMB2 (Abcam, Cambridge, UK), anti-CXCR4 clone 2B11 (Invitrogen), and the appropriate secondary antibodies coupled to Alexa 488 (Invitrogen) or Alexa 568 (Abcam). Slides were imaged on a Nikon Eclipse Ti2 immunofluorescence microscope (Nikon, Tokyo, Japan).

### Flow Cytometry

Cells/engrafted organoids were treated with PAMD-Ch17 at the indicated dose for the indicated time. Viability was determined by 7-amino-actinomycin D (7-AAD) (Biolegend, San Diego, CA) or Live/Dead Fixable Blue (Invitrogen) staining. Mitochondrial superoxide was measured by MitoSOX Red (Invitrogen). Mitochondrial membrane potential was measured by MitoProbe TMRM (Thermo Scientific). Surface CD8 levels were measured by staining with the anti-CD8 antibody, clone 53-6.7 (Biolegend). Surface CXCR4 levels were measured by staining with the anti-CXCR4 antibody, clone 12G5 (BioLegend). Surface CXCR4 blocking was measured by staining with clone 12G5 (PE) and clone 2B11 (eFluor450) (Invitrogen). For fPAMD uptake, cells were treated with 0.6uM fPAMD for 30 minutes. Data were collected on the BD LSRII (BD Biosciences, Franklin Lakes, NJ). Data were then analyzed using FlowJo software v10.7.0 (Treestar, Ashland, OR).

### Migration Assay

Migration assay was performed using the 3um chemotaxis assay 24 well format kit (Cell Biolabs, San Diego, CA) as per manufacturer’s instructions and CXCL12 (Peprotech, Cranbury, NJ). Migration was read using a Tecan Infinite M200 PRO in fluorescence top reading mode with excitation 480nm and emission 520nm. Readings were normalized to the control condition.

### Seahorse Assay

Cells were treated with PAMD-Ch17 at the indicated dose for the indicated time, counted, and equal numbers of viable cells plated per condition into Seahorse XF RMPI medium pH 7.4 (Aglient, Santa Clara, CA) supplemented with 0.01M glucose, 1mM pyruvate, and 2mM L-glutamine. Cell counts were checked using a CeligoImage Cytometer (Celigo, Redwood City, CA). Cells were then processed for seahorse assay as per manufacturer’s instructions.

### Cloning and Lentiviral Transduction

*CXCR4* KO Jurkat cells were generated using the guide RNAs (gRNAs) in ^27^. gRNAs were cloned into LentiCRISPRv2GFP plasmid (Addgene plasmid # 82416) and transformed into Stellar Competent Cells (Takara) ^28,29^. Cells were plated, single colonies obtained, and plasmids sequenced. A validated plasmid for each gRNA was subsequently used. Lenti-X 293T cells were transfected with second generation lentiviral plasmids and the gRNA plasmid using Lipofectamine 2000 (Invitrogen) according to manufacturer’s protocol. Cells were transduced with 10ug/ml polybrene (EMD Millipore, Burlington, MA). GFP^+^ cells were sorted into single cells in a 96 well plate to obtain single cell subclones.

### Statistical Analysis

Statistical testing was performed using GraphPad Prism v10.0.2 (Graphpad Software, La Jolla, CA). We used students t-test and one-way ANOVA as appropriate. EC50s were calculated using log(inhibitor) vs response variable slope four parameters fitting. We display mean±standard error of mean. P≤0.05 was considered significant. Sample size given in the text.

## RESULTS

### PAMD-Ch17 is cytotoxic against a variety of human acute leukemias, including T cell acute lymphoblastic leukemia

To begin investigating the mechanism of PAMD-Ch17’s anti-leukemic effects, we performed MTT assays on a panel of human AML and ALL cell lines. Cells were treated with increasing doses of PAMD-Ch17 for 72 hours, analyzed, and the effective concentrations at which cells showed a 50% decrease in metabolic activity (EC50) was calculated for each cell line. We found that PAMD-Ch17 reduced metabolic activity against all cell lines tested, with B-ALL and T-ALL cell lines showing a lower EC50 as compared to AML cell lines (Figure 1A and B). We confirmed that PAMD-Ch17’s parent molecule, AMD3100, did not affect metabolic activity at even the highest dose tested (5µM), consistent with previous studies (Supplemental Figure 1) ^15,17,30-33^.

**Figure 1.**
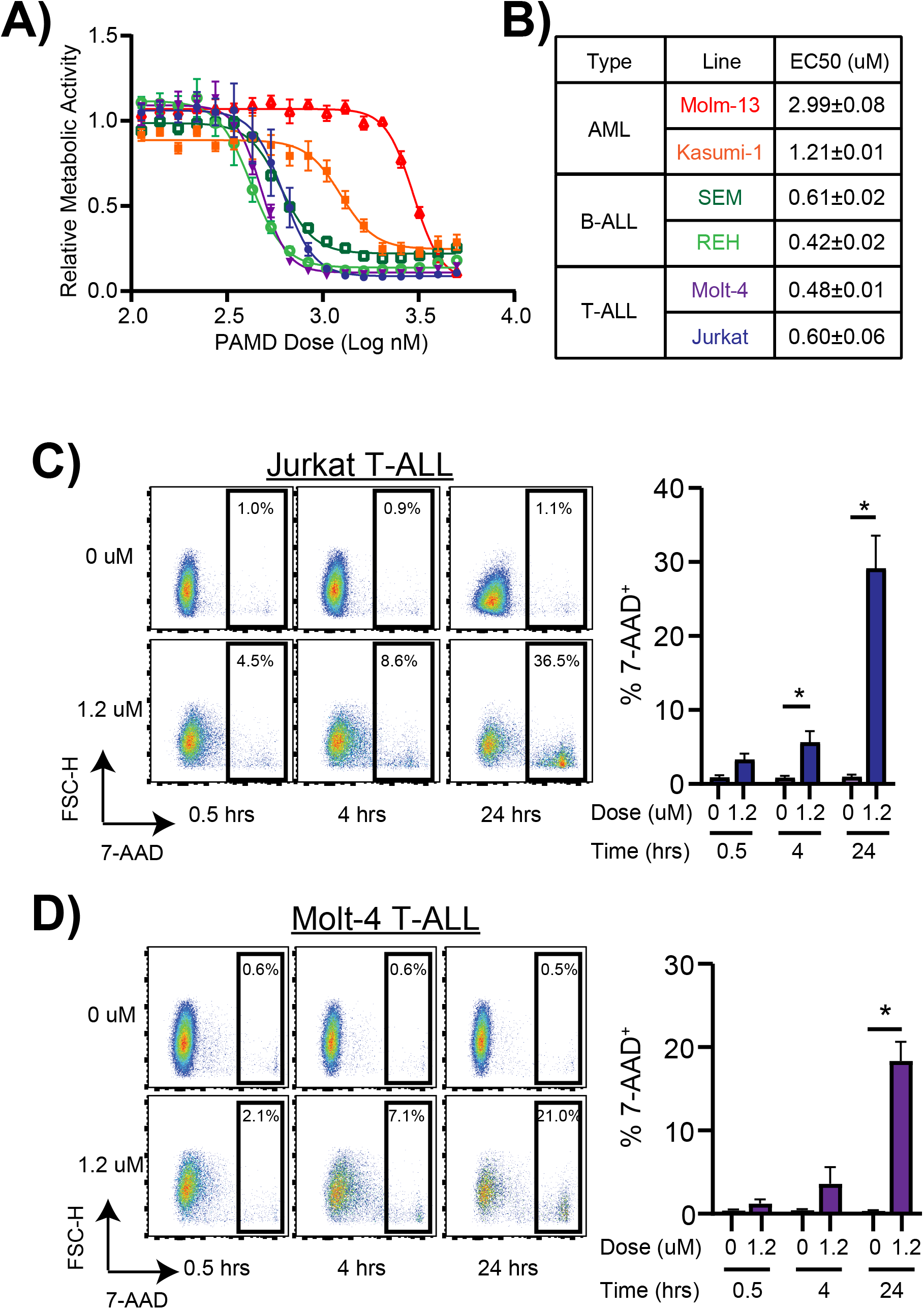
PAMD-Ch17 induces cytotoxicity in human acute leukemia cell lines. A) Dose-response curves of relative metabolic activity for the indicated human AML and ALL cell lines treated with PAMD-Ch17 for 72 hours. B) Table with colors denoting the cell line, leukemia type, and EC50s shown in A. C) Representative flow cytometry plots and bar graph of 7-AAD staining in Jurkat cells and (D) Molt-4 cells treated with PAMD-Ch17 for the indicated dose and time. N=3. * p ≤ 0.05.

We previously showed that PAMD-Ch17 induces cell death in AML cells ^13^. Given the lower EC50s of the T-ALL cell lines, we decided to focus this study on T-ALL. To test if PAMD-Ch17 similarly affects the viability of T-ALL cells, we assayed cell viability in Jurkat and Molt-4 cells. We treated cells with vehicle or 1.2 µM of PAMD-Ch17 for 0.5, 4, or 24 hours, assayed viability using 7-AAD and analyzed by flow cytometry. In both Jurkat and Molt-4 cells, PAMD-Ch17 induced a notable, statistically significant increase in cell death with the 1.2µM dose, starting at 4 hours and strongly at 24 hours (Figure 1C and D). These results indicate that PAMD-Ch17 has anti-leukemic effects against human T-ALL cells, and at lower doses than with AML cell lines.

### PAMD-Ch17 inhibits CXCR4 but does not require the cytokine receptor for its anti-leukemic effects

PAMD-Ch17’s only known target is the CXCR4 receptor, raising the possibility that PAMD-Ch17 mediates its anti-leukemic effects through the receptor ^6-14^. To test this, we first confirmed that PAMD-Ch17 inhibits CXCR4 in leukemia cells. Jurkat T-ALL cells were treated with either 0.6µM of the CXCR4 inhibitor AMD3100 or PAMD-Ch17, then stained with two different antibodies against CXCR4 and analyzed by flow cytometry ^34-36^. Clone 12G5 binds to the CXCL12 binding site of CXCR4, the CXCR4 binding site of AMD3100 and the expected binding site of PAMD-Ch17 ^34,35^. The second antibody, 2B11, binds to the N-terminus of CXCR4 and is not expected to interfere with either AMD3100 or PAMD-Ch17 binding to CXCR4 ^36^. We found that PAMD-Ch17, like AMD3100, decreased staining with the CXCR4 12G5 antibody, implying that the polymer blocks the CXCL12 binding site on CXCR4 (Figure 2A, Supplemental Figure 2). Notably, PAMD-Ch17 significantly decreased 12G5 staining more than AMD3100 did. Neither PAMD-Ch17 nor AMD3100 blocked binding of the N-terminal antibody, indicating that cell surface expression of the receptor is not affected (Figure 2B). We then tested if PAMD-Ch17 blocks CXCR4 function using migration assays ^6-14^. Jurkat cells were treated with either 2.5ug/ml of PAMD-Ch17 or 0.3uM of AMD3100 as in previous studies, then allowed to migrate in response to 20nM CXCL12, the canonical ligand for CXCR4 ^12,37^. PAMD-Ch17 showed a trend towards decreased migration similar to AMD3100, supporting that PAMD-Ch17 functionally inhibits CXCR4 (Figure 2C).

**Figure 2.**
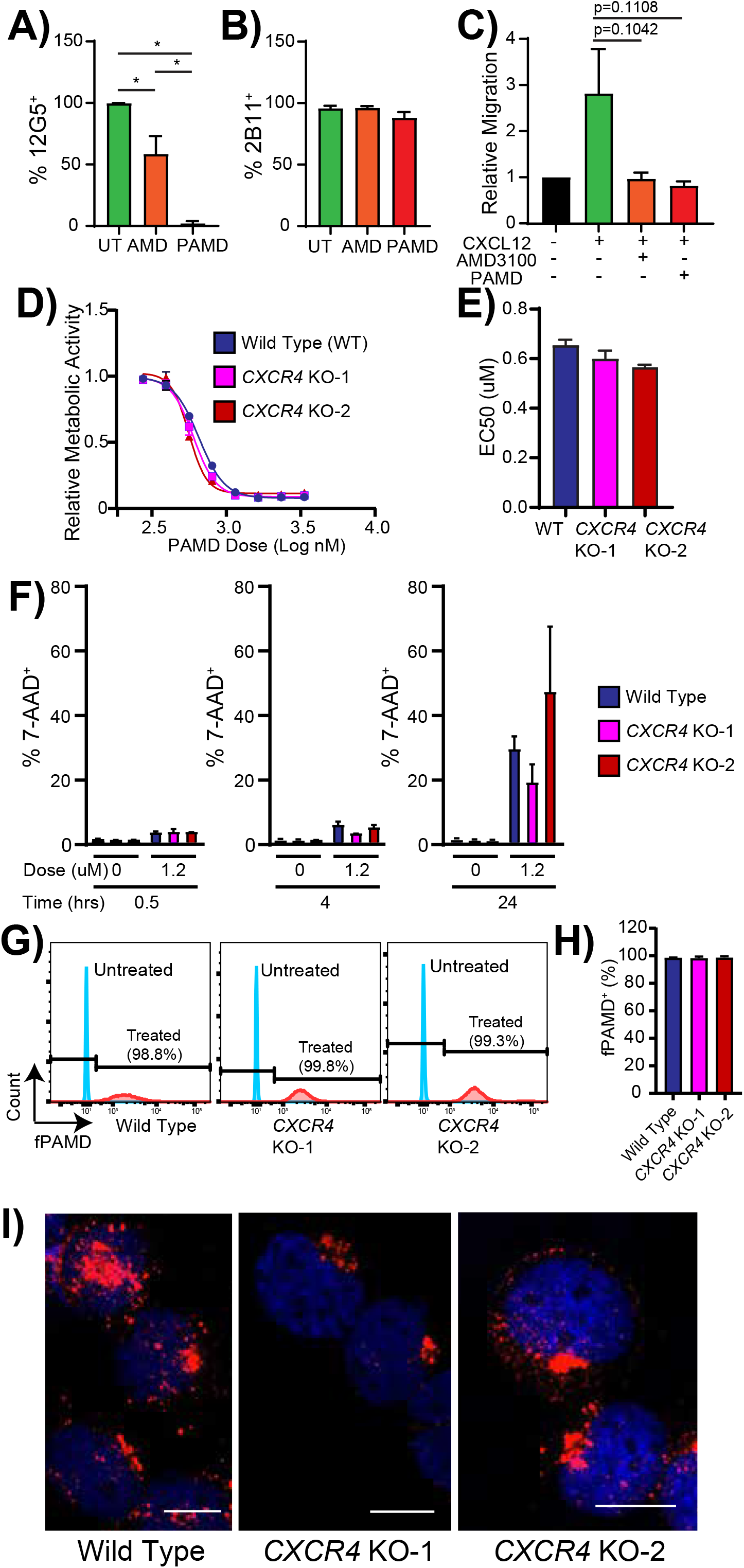
PAMD-Ch17 inhibits the CXCR4 receptor but does not require CXCR4 for its anti-leukemic effects. A) Graph of percentage (%) of Jurkat cells recognized by the anti-CXCR4 12G5 or B) 2B11 antibody after treatment for 24 hours with either 0.6µM AMD3100, PAMD-Ch17, or untreated. C) Graph of relative migration in response to CXCL12 of Jurkat cells treated with either AMD3100, PAMD-Ch17, or untreated for 3 hours. D) Dose-response curves of relative metabolic activity for wild type and *CXCR4* KO Jurkat cells, as measured by MTT assay. E) Graph of the EC50s of the indicated cells treated as in D. F) Graph of the % 7-AAD stained Jurkat cells of the indicated genotype treated with 0 µM or 1.2 µM of PAMD-Ch17 for 0.5, 4, or 24 hours. G) Representative flow cytometry plots of Jurkat cells of the indicated genotype treated with a fluorescently labelled PAMD-Ch17 (fPAMD) or untreated. H) Graph of the % of fluorescent cells of the indicated genotype after treatment as in G. I) Confocal microscopy images of cells treated as in G. N=3. * p ≤ 0.05.

We then tested if PAMD-Ch17 requires CXCR4 to mediate its anti-leukemic effects. To do so, we generated *CXCR4* knock out (KO) Jurkat cells using Crispr/Cas 9 ^27-29^. We obtained two *CXCR4* KO subclones, hereafter referred to as *CXCR4* KO-1 and *CXCR4* KO-2. We validated that both KO clones lost CXCR4 expression by flow cytometry and microscopy (Supplemental Figure 3A and B) ^34-36^. We confirmed loss of CXCR4 activity using migration assays (Supplemental Fig.3C) ^12,37^. Altogether, these data confirm that we successfully knocked out CXCR4 in both Jurkat subclones.

We then tested the effect of PAMD-Ch17 on the *CXCR4* KO cells. By MTT assay, we found that PAMD-Ch17 had similar effects on metabolic activity and EC50s against *CXCR4* KO Jurkat and wildtype cells (Figure 2D and E). By 7-AAD staining, we found that PAMD-Ch17 induced similar levels of cell death in *CXCR4* KO leukemia cells as in wildtype cells (Figure 3F). To test if *CXCR4* KO cells still uptake the drug, we used a fluorescently labelled PAMD-Ch17 (fPAMD) and found that *CXCR4* KO leukemia cells internalized the drug similarly to the wild type cells (Figure 2G and H). Curiously, in these cells, fPAMD localized to the perinuclear region (Figure 2I). These data indicate that while PAMD-Ch17 inhibits CXCR4, PAMD-Ch17 uptake and its anti-leukemic activity are independent of CXCR4.

**Figure 3.**
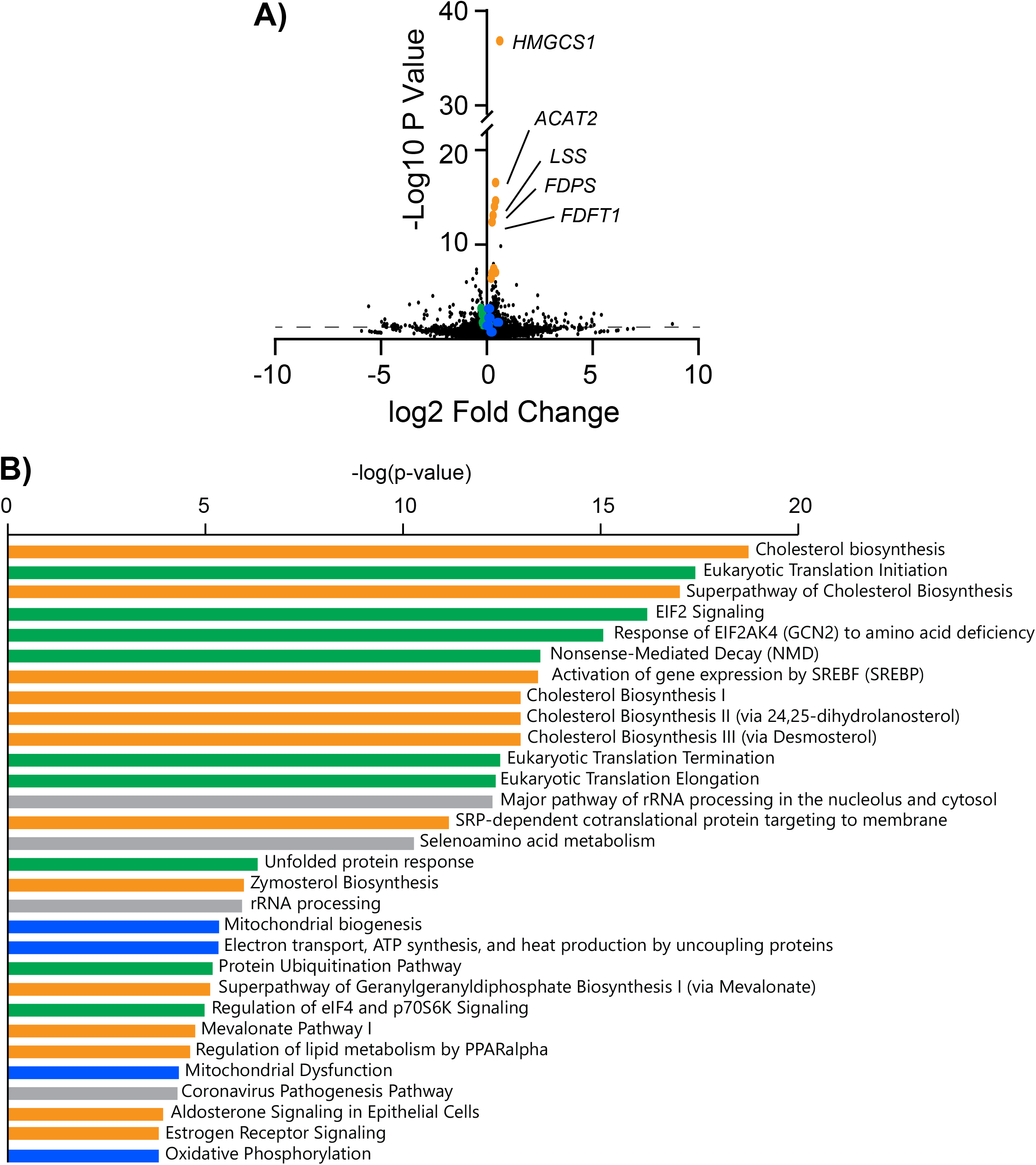
PAMD-Ch17 induces differential expression of genes related to cholesterol biosynthesis, protein translation, and mitochondrial homeostasis. A) Volcano plot of differentially expressed genes in Jurkat cells treated with 0.6uM PAMD-Ch17 for 24 hours as compared to untreated cells. Selected cholesterol related genes are colored in yellow, translation in green, and mitochondria in blue. B) Top 30 pathways associated with the differentially associated genes identified using Ingenuity Pathway Analysis. Orange color denotes cholesterol related pathways, green is translation pathways, blue mitochondrial related pathways, and gray are other pathways.

### PAMD-Ch17 dysregulates expression of genes involved in cholesterol biosynthesis, translation, and mitochondria function

To gain broad mechanistic insight into PAMD-Ch17’s activity, we performed whole transcriptome sequencing on untreated and PAMD-Ch17 treated Jurkat cells. Jurkat cells were treated with 0.6µM PAMD-Ch17 or vehicle for 24 hours, RNA was isolated, libraries prepared, reads filtered, and differentially expressed genes identified ^22-25^. PAMD-Ch17 induced differential expression of 1269 genes, with 694 genes downregulated and 575 genes upregulated (Figure 3A). By Ingenuity Pathway Analysis, we found that the genes dysregulated by PAMD-Ch17 are involved in numerous cellular pathways, including cholesterol synthesis, protein translation, and mitochondrial homeostasis (Figure 3B) ^22-25^. A list of the differentially expressed genes can be found in Supplemental Table 1.

To determine whether these changes in gene expression are due to PAMD-Ch17’s CXCR4 independent activities, we performed whole transcriptome sequencing on *CXCR4* KO cells. We found that loss of CXCR4 induced differential expression of 1648 genes, with only 78 (<10%) shared with PAMD-Ch17 treated Jurkat cells. (Supplemental Figure 4, Supplemental Table 1). This indicates that the majority of the gene expression changes induced by PAMD-Ch17 are related to its CXCR4 independent activities.

### PAMD-Ch17 colocalizes with the mitochondria in leukemia cells

Because PAMD-Ch17 induced differential expression of genes related to mitochondrial function and accumulates in the perinuclear region where mitochondria typically reside, we asked whether PAMD-Ch17 mediates its anti-leukemic effects through mitochondrial dysfunction. To address this possibility, we tested if PAMD-Ch17 colocalizes with the mitochondria. We treated Jurkat cells with fPAMD at the indicated dose and length of time ^13^. Cells were harvested, affixed to glass slides by cytospin and mitochondria were stained with Mitotracker Deep Red and co-localization was determined using Manders Colocalization Coefficient M1 and M2 ^26,38^. At 4 hours, there was moderate co-localization of the drug at the mitochondria while at 24 hours there was strong co-localization by Manders Colocalization Coefficient M1, suggesting that the majority of the drug accumulated in the mitochondria (Figure 4A and B, Supplemental Figure 5). Co-localization of the mitochondria to the drug signal, Manders Colocalization Coefficient M2, remained moderate and constant, suggesting that at the dose tested, not all mitochondria accumulated drug (Figure 4C and Supplemental Figure 5).

**Figure 4.**
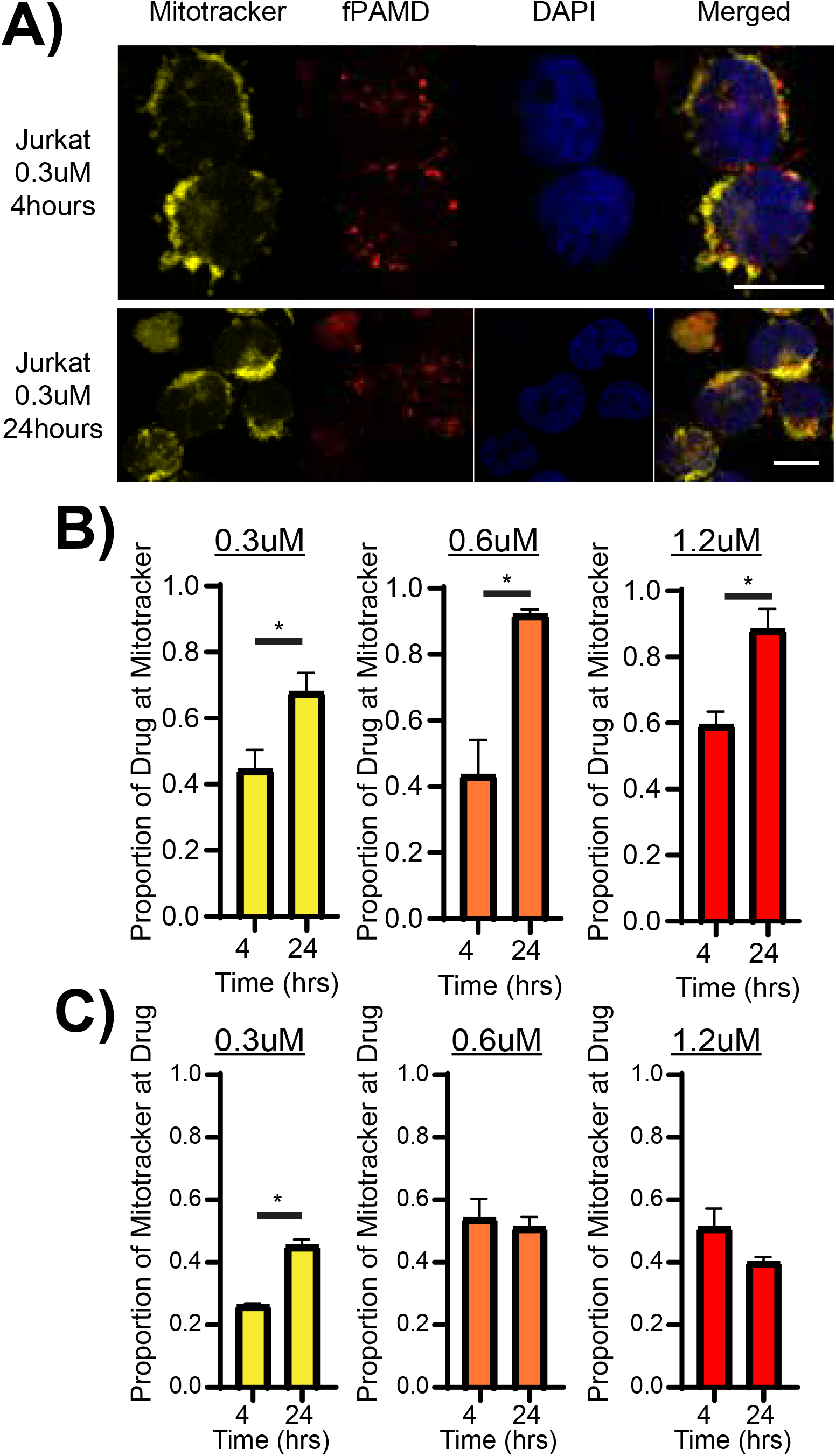
PAMD-Ch17 colocalizes with the mitochondria. A) Representative confocal images of Jurkat cells treated with 0.3uM fluorescently tagged PAMD-Ch17 (fPAMD) for 4 or 24 hours, then stained with Mitotracker Deep Red and DAPI. B) Graph of Mitotracker Deep Red overlap at fPAMD at the indicated dose and time by Mander’s Colocalization Coefficient M1. C) Graph of fPAMD overlap at Mitotracker Deep Red at the indicated dose and time by Mander’s Colocalization Coefficient M2. N=3. Scale bar is 10um. * p ≤ 0.05.

### PAMD-Ch17 induces mitochondrial superoxide, loss of membrane potential, and decreases mitochondrial function

To test if PAMD-Ch17 induces mitochondrial dysfunction, we tested whether PAMD-Ch17 induces mitochondrial superoxide, a hallmark of damaged mitochondria ^39-50^. Jurkat and Molt-4 cells were treated with 0µM or 1.2µM PAMD-Ch17 for 0.5, 4, or 24 hours, and mitochondrial superoxide assayed using MitoSox Red. We found that PAMD-Ch17 induced mitochondrial superoxide starting at 0.5 hours of treatment in a time dependent manner (Figure 5A and C). To further test PAMD-Ch17’s effect on mitochondrial function, we assayed the polymer’s effect on mitochondria membrane potential. Jurkat and Molt-4 cells were treated with 0µM or 1.2µM PAMD-Ch17 for 0.5, 4, or 24 hours and stained with Tetramethylrhodamine methyl ester (TMRM), a marker of mitochondrial membrane potential integrity. PAMD-Ch17 decreased the percentage of cells with intact membrane potential starting at 0.5 hours in Jurkat cells and at 4 hours in Molt 4 cells (Figure 5B and D).

**Figure 5.**
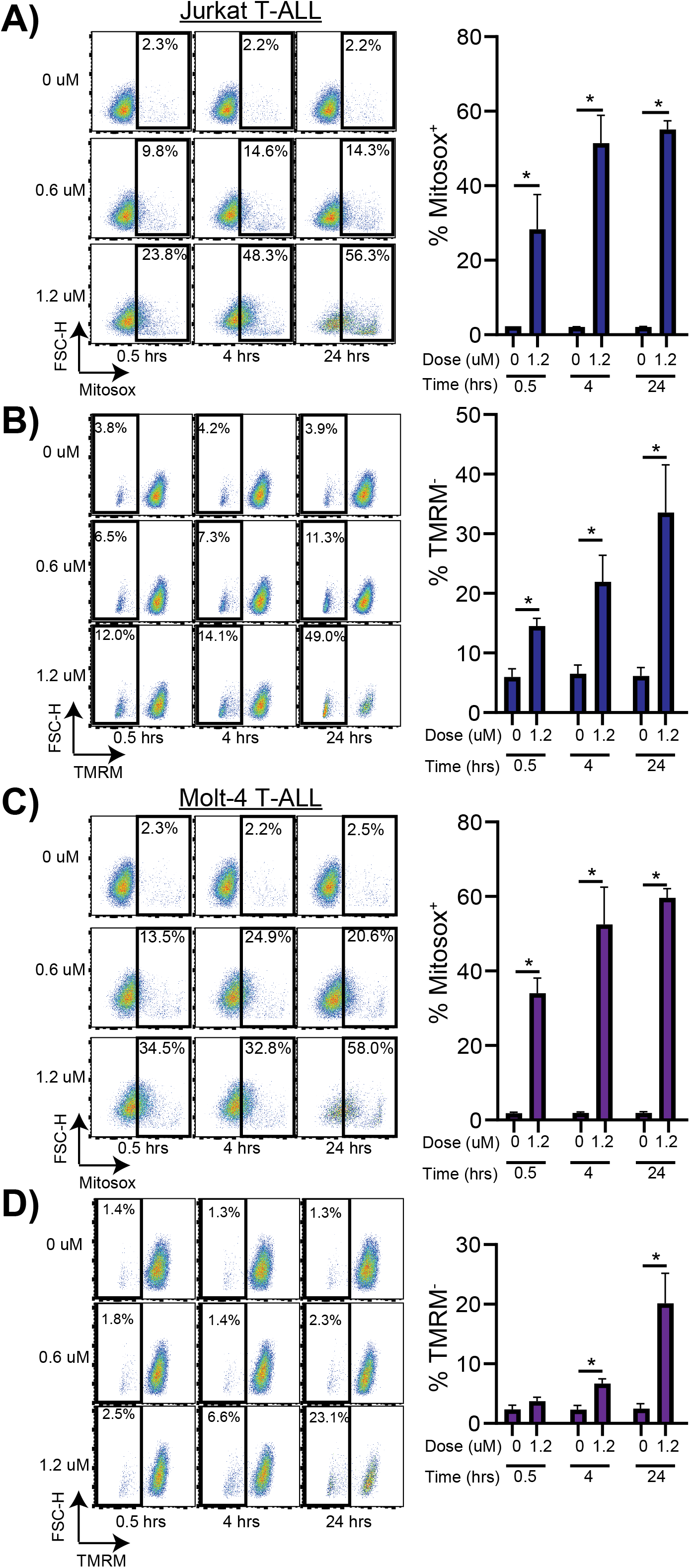
PAMD-Ch17 induces mitochondrial superoxide and loss of mitochondrial membrane potential. A) Representative flow cytometry plots of Jurkat cells treated with 0 µM or 1.2 µM PAMD-Ch17 for 0.5, 4, or 24 hours, then stained for mitochondrial superoxide with Mitosox Red. Graph of the percentage (%) of Mitosox positive cells is shown on the right. B) Representative flow cytometry plots of Jurkat cells treated as in A) and stained for mitochondrial membrane potential with Tetramethylrhodamine methyl ester (TMRM). Graph of the percentage (%) of TMRM negative cells is shown on the right. C) Representative flow cytometry plots of Molt-4 cells treated and stained as in A) and D) as in B). N=3. * p ≤ 0.05.

To test if PAMD-Ch17 impairs mitochondrial metabolism, we performed Seahorse assays. Jurkat T-ALL cells were treated with 0µM or 1.2µM PAMD-Ch17 for 24 hours and equal numbers of viable cells plated for analysis. PAMD-Ch17 decreased mitochondrial respiration overall (Figure 6A) and decreased basal oxygen consumption, oxygen linked ATP production, and proton leakage (Figure 6B). Non-mitochondrial oxygen consumption and maximal oxygen consumption were not significantly changed (Figure 6C). Analysis of the extracellular acidification rate revealed that PAMD-Ch17 also decreased baseline non-mitochondrial metabolism overall (Figure 6D and E). Collectively, these results indicate that PAMD-Ch17 induces significant mitochondrial dysfunction, providing a potential explanation for its anti-leukemic activity.

**Figure 6.**
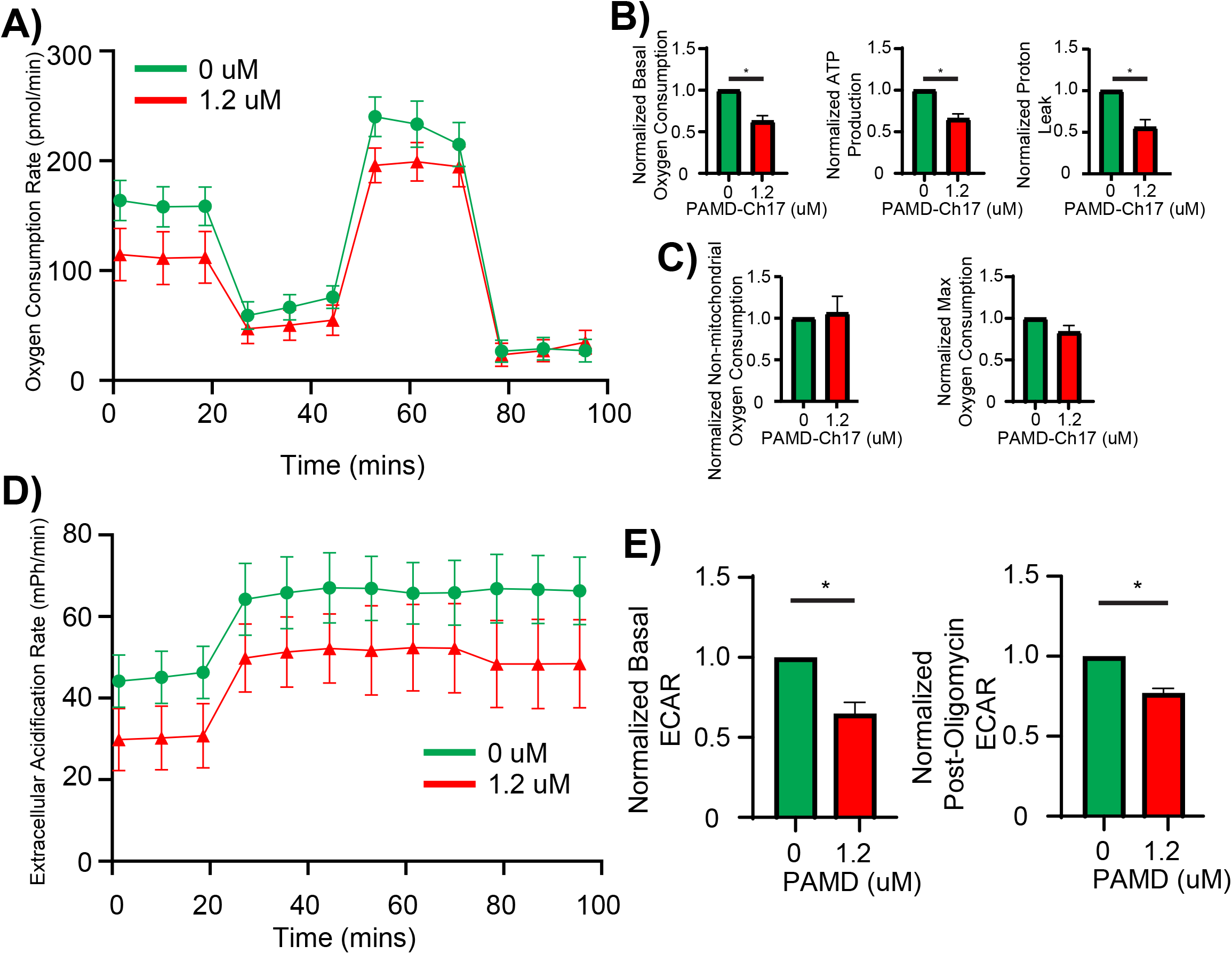
PAMD-Ch17 induces mitochondrial dysfunction. A) Representative graph of the Oxygen Consumption Rate for Jurkat cells treated with the indicated dose of PAMD-Ch17 for 24 hours. B) Graphs of normalized basal oxygen consumption, ATP production, and proton leakage from the mitochondria in cells treated as in A). C) Graphs of normalized non-mitochondrial oxygen consumption and maximum (max) oxygen consumption in cells treated as in A). D) Representative graph of the Extracellular Acidification Rate (ECAR) and E) graphs of normalized basal and post-oligomycin ECAR from Jurkat cells treated as in A. N=3. * p ≤ 0.05.

### PAMD-Ch17 induces mitochondrial superoxide and cell death in primary T-ALL cells

To determine if PAMD-Ch17 causes mitochondrial dysfunction and cell death specifically in primary T-ALL, we deployed two different models: a viral expression mouse model of T-ALL, and a human bone marrow organoid model. To generate mouse T-ALL cells we transduced bone marrow cells with virus containing the constitutively active ΔEGFΔLNRΔP Notch1 mutant receptor and GFP from an internal ribosomal entry site ^18,19^. Transduced cells were transplanted into congenic mice. One to two months after transplantation, mice developed a GFP^+^, CD8^+^ leukemia as previously reported for this model (Supplemental Figure 6A) ^18^. Leukemic mice were then sacrificed on first sign of distress, bone marrow harvested, and cells treated *in vitro* with 0µM (UT) or 1.2µM PAMD-Ch17 (T) for 4 or 24 hours. Bone marrow from healthy mice was similarly harvested and treated with PAMD-Ch17. Cells were then stained for either mitochondrial superoxide using MitoSox Red or viability using 7-AAD. PAMD-Ch17 significantly induced mitochondrial superoxide and cell death at both time points in GFP^+^ mouse leukemia cells (Figure 7A and B). PAMD-Ch17 also induced mitochondrial superoxide and cell death in healthy bone marrow cells, but at lower levels than in T-ALL cells. This difference in cell death between leukemia and healthy cells was particularly striking, consistent with previous results (Supplemental Figure 6B and C) ^13^.

**Figure 7.**
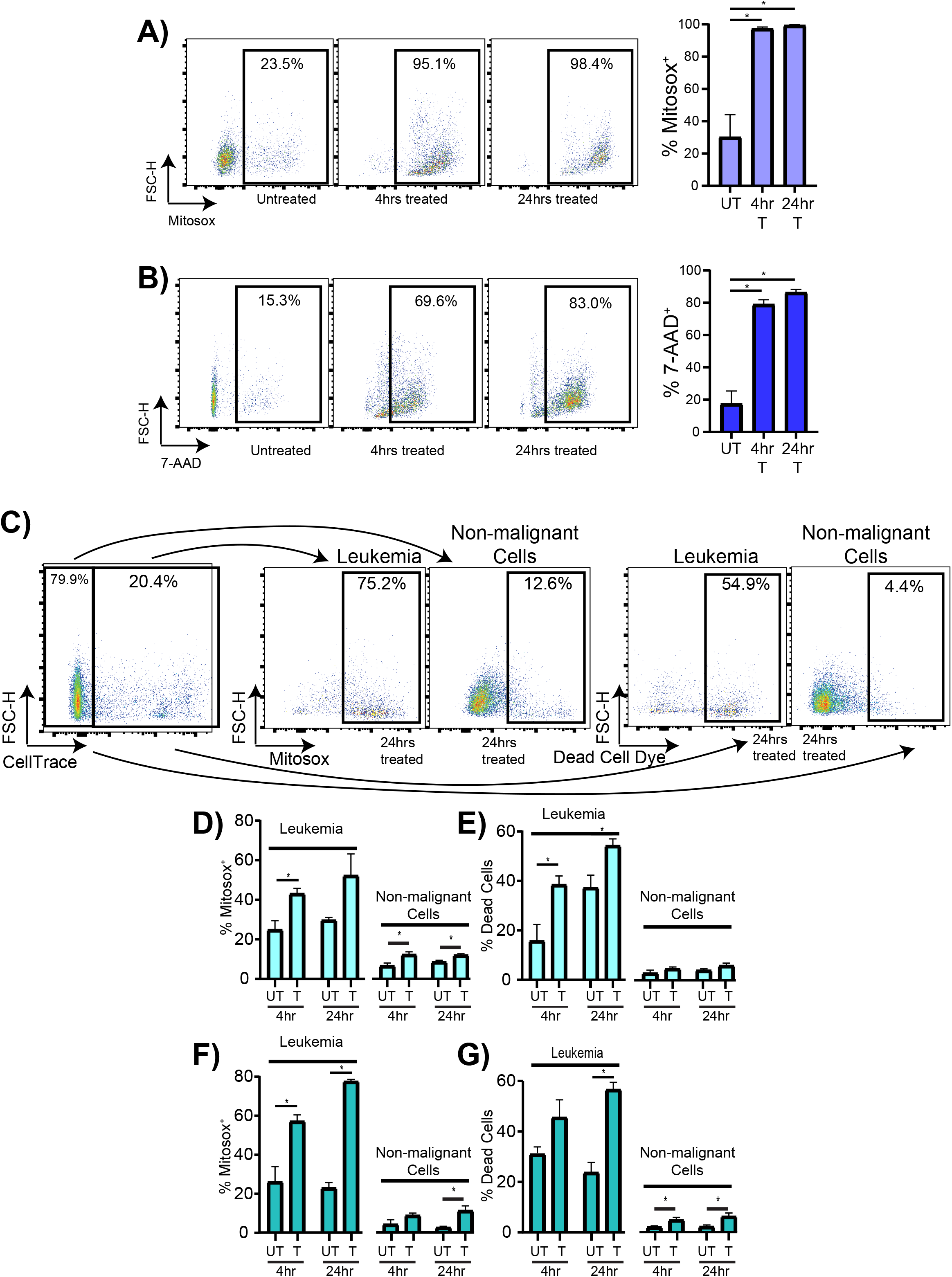
PAMD-Ch17 induces mitochondrial superoxide and cell death in primary T-ALL cells, with minimal effects on healthy cells. Mouse primary T-ALL cells generated by viral overexpression of the hyperactive Notch 1 ΔEGFΔLNRΔP construct were treated with 1.2uM of PAMD-Ch17 *in vitro* for the indicated time and stained for mitochondrial superoxide with Mitosox Red and viability using 7-AAD. A) Representative flow cytometry plots and graph of the percentage (%) of Mitosox positive (+) and B) 7AAD^+^ mouse Notch 1 ΔEGFΔLNRΔP expressing T-ALL cells. N=4 mouse primary T-ALL samples. C) Representative flow cytometry plots from human bone marrow (BM) organoids engrafted with CellTrace labeled human primary T-ALL cells and treated with 0.6uM PAMD-Ch17 for the indicated time. D) Graph of the percentage (%) of Mitosox^+^ and E) dead T-ALL cells and non-malignant cells from organoids engrafted with patient sample #1. F, G) Same as D and E from organoids engrafted with patient sample #2. UT=untreated. T=PAMD-Ch17 treated. 5-7 engrafted organoids were combined for each replicate, and 3 replicates for each human primary T-ALL sample were used for each condition. * p ≤ 0.05.

To test the effect of PAMD-Ch17 in a human system, we used bone marrow organoids ^20,21^. Human induced pluripotent stem cells were induced to differentiate into the stromal and hematopoietic cells of the bone marrow and then engrafted with primary T-ALL cells from two different patients ^20,21^. Three-four days after engraftment, organoids were treated with 0µM (UT) or 0.6µM PAMD-Ch17 (T) for 4 or 24 hours and stained for mitochondrial superoxide using MitoSox Red or viability using Live/Dead Fixable Blue. Using the Cell Trace dye to differentiate the T-ALL cells from the non-malignant cells, we found that leukemia cells made up 10-20% of the total cells in the organoids (Figure 7C, Supplemental Figure 7). We found that PAMD-Ch17 induced higher levels of mitochondrial superoxide in the engrafted T-ALL cells as compared to the non-malignant cells in the organoids (Figure 7D and F). PAMD-Ch17 also induced cell death at higher levels in the leukemia cells compared to the healthy cells (Figure 7E and G). These findings show that PAMD-Ch17 induces mitochondrial dysfunction and cell death in primary T-ALL at markedly higher levels than in healthy bone marrow cells, providing a potential mechanism for the polymer’s anti-leukemic effects.

## DISCUSSION

In our previous work, we showed that the polymeric drug PAMD-Ch17 has novel anti-leukemic effects against AML that are not shared with other CXCR4 inhibitors ^13^. In this study, we show that PAMD-Ch17 is also effective against T-ALL and, although it effectively inhibits CXCR4, the polymeric drug does not mediate its anti-leukemic effects through the receptor. Our whole transcriptome sequencing data suggested mitochondrial involvement. In fact, we found that PAMD-Ch17 colocalizes to the mitochondria and induces mitochondrial stress, damage and dysfunction in human T-ALL cell lines. In addition, we found that PAMD-Ch17 induces mitochondrial superoxide and cell death in both mouse primary T-ALL and human primary T-ALL cells, but not in healthy mouse bone marrow or non-malignant human bone marrow organoid cells. Collectively, our results provide a possible mechanism for PAMD-Ch17’s novel anti-leukemic effects, an explanation for its selectivity, and support the development of this class of drugs for the treatment of leukemia.

Our data suggests that PAMD-Ch17 directly targets the mitochondria. Specifically, we observed that PAMD-Ch17 co-localized at the mitochondria within 4 hours, induced significant mitochondrial superoxide starting at 0.5 hours, and significantly decreased the percentage of cells with intact membrane potential between 0.5 and 4 hours, depending on the cell line tested. In contrast, we did not observe a significant increase in cell death until 4-24 hours. These data indicate that the induced mitochondrial dysfunction is more likely a cause, rather than the consequence of cell death. Leukemias are known to have increased reliance on mitochondria respiration as compared to healthy hematopoietic cells ^39-57^. This increased mitochondrial reliance is associated with increased baseline reactive oxygen species (ROS) levels. As a result, cancer cells are more sensitive to any further increases in ROS, as compared to healthy cells ^39,40,46-54^. In addition, leukemia cells are also known to have physically aberrant mitochondria, with different protein expression, DNA content, morphology, and functional capacity ^42-45^. These differences could explain why PAMD-Ch17 effectively kills leukemia cells, while having minimal effects on healthy cells.

While we have identified a potential mechanism of action for PAMD-Ch17, we cannot conclude that the mitochondria are the polymer’s sole target in leukemia cells. Our co-localization analysis indicates that the drug may be localizing to other regions of the cell, especially at the 4 hour timepoint. In addition, our whole transcriptome sequencing data indicate that PAMD-Ch17 altered expression of genes involved in cholesterol biosynthesis and translation which are both hallmarks of endoplasmic reticulum (ER) dysfunction. It is possible that the mitochondrial impairment caused by PAMD-Ch17 leads to ER dysfunction. However, it is equally possible that PAMD-Ch17 disrupts the activity both organelles ^58-63^.

It remains unknown how PAMD-Ch17 enters leukemia cells, as CXCR4 is not required for drug uptake. Other CXCR4 related receptors such as CCR7, CXCR7, and CD74 were not found to be expressed on Jurkat cells (data not shown), and are thus unlikely to be mediating uptake ^64^. Uptake of related PAMD variants with fluorescently labelled siRNAs in CXCR4^+^ non-leukemia cell lines yield patterns similar to that observed in our T-ALL cells: distinct puncta with notable perinuclear localization ^6,9,14^. The lack of diffuse uptake and the formation of distinct puncta hint at an active uptake mechanism and that the uptake mechanism may be similar across different cell types. It will require further investigation to determine whether the uptake mechanism is receptor dependent, cell type specific, and/or through an active process.

Our data lead us to propose the following mechanism: PAMD-Ch17 localizes to the mitochondria to induce mitochondrial superoxide that damages the mitochondria and its function, thereby leading to leukemia cell death (See Graphical Abstract). PAMD-Ch17’s novel, selective anti-leukemic effects support its potential as a therapeutic agent. In conjunction with its potential to be a chemosensitizer via CXCR4 inhibition, and ability form nanoparticle nucleic acid delivery vehicles, PAMD-Ch17 could be a potent multi-function therapeutic. Our results provide insight towards the development of PAMDs as anti-leukemic agents.

## ACKNOWLEDGEMENTS

We thank Dr. Rebecca Oberly-Deegan, Dr. Micah Schott, and Dr. Amar Natarajan along with their lab members for technical assistance and the equipment required for this study. We thank Dr. Sarah Holstein, Dr Sidharth Mahapatra, and Dr. Joshi for their feedback on this project. We thank Dr. Warren Pear for providing us with the ΔEGFΔLNRΔP construct and advice for generating the mouse T-ALL model. We thank the UNMC Flow Cytometry Core, UNMC Sequencing Core, and UNMC Microscopy Core.

## AUTHOR CONTRIBUTIONS

C.L., S.R., D.O., R.K.H. conceived of and designed the study. C.J., E.K., S.T., S.R., D.O., provided PAMD-Ch17 for experiments. C.L., A.D., S.P., V.R., E.M., A.R.B., S.S. D.C., K.H., performed experiments. X.P. analyzed whole transcriptome sequencing data. C.L., D.O., R.K.H. wrote the manuscript.

## Graphical Abstract

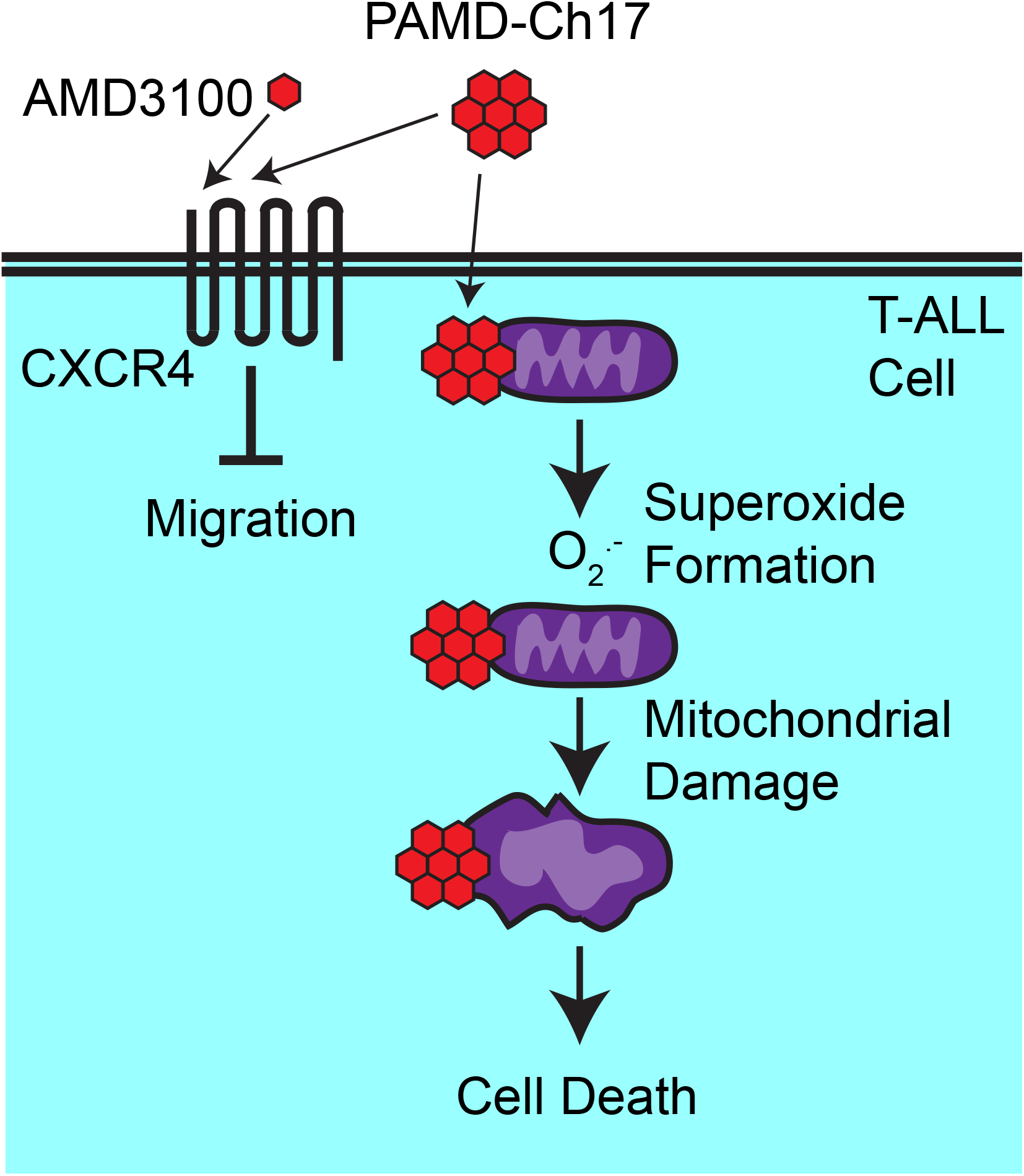

## REFERENCES

1. Raetz EA, Teachey DT. T-cell acute lymphoblastic leukemia. Hematology Am Soc Hematol Educ Program. 2016;2016(1):580–588.

2. Litzow MR, Ferrando AA. How I treat T-cell acute lymphoblastic leukemia in adults. Blood. 2015;126(7):833–841.

3. Marks DI, Paietta EM, Moorman AV, et al. T-cell acute lymphoblastic leukemia in adults: clinical features, immunophenotype, cytogenetics, and outcome from the large randomized prospective trial (UKALL XII/ECOG 2993). Blood. 2009;114(25):5136–5145.

4. Marks DI, Rowntree C. Management of adults with T-cell lymphoblastic leukemia. Blood. 2017;129(9):1134–1142.

5. Douvas MG, Riegler LL. Meeting Challenges in the Long-Term Care of Children, Adolescents, and Young Adults with Acute Lymphoblastic Leukemia. Curr Hematol Malig Rep. 2022;17(1):15–24.

6. Hang Y, Tang S, Tang W, et al. Polycation fluorination improves intraperitoneal siRNA delivery in metastatic pancreatic cancer. J Control Release. 2021;333:139–150.

7. Li J, Oupicky D. Effect of biodegradability on CXCR4 antagonism, transfection efficacy and antimetastatic activity of polymeric Plerixafor. Biomaterials. 2014;35(21):5572–5579.

8. Li J, Zhu Y, Hazeldine ST, Li C, Oupicky D. Dual-function CXCR4 antagonist polyplexes to deliver gene therapy and inhibit cancer cell invasion. Angew Chem Int Ed Engl. 2012;51(35):8740–8743.

9. Tang S, Kapoor E, Ding L, et al. Effect of tocopherol conjugation on polycation-mediated siRNA delivery to orthotopic pancreatic tumors. Biomater Adv. 2023;145:213236.

10. Wang Y, Kumar S, Rachagani S, et al. Polyplex-mediated inhibition of chemokine receptor CXCR4 and chromatin-remodeling enzyme NCOA3 impedes pancreatic cancer progression and metastasis. Biomaterials. 2016;101:108–120.

11. Wang Y, Li J, Chen Y, Oupicky D. Balancing polymer hydrophobicity for ligand presentation and siRNA delivery in dual function CXCR4 inhibiting polyplexes. Biomater Sci. 2015;3(7):1114–1123.

12. Wang Y, Li J, Oupicky D. Polymeric Plerixafor: effect of PEGylation on CXCR4 antagonism, cancer cell invasion, and DNA transfection. Pharm Res. 2014;31(12):3538–3548.

13. Wang Y, Xie Y, Williams J, et al. Use of polymeric CXCR4 inhibitors as siRNA delivery vehicles for the treatment of acute myeloid leukemia. Cancer Gene Ther. 2020;27(1-2):45-55.

14. Xie Y, Wehrkamp CJ, Li J, et al. Delivery of miR-200c Mimic with Poly(amido amine) CXCR4 Antagonists for Combined Inhibition of Cholangiocarcinoma Cell Invasiveness. Mol Pharm. 2016;13(3):1073–1080.

15. Uy GL, Rettig MP, Motabi IH, et al. A phase 1/2 study of chemosensitization with the CXCR4 antagonist plerixafor in relapsed or refractory acute myeloid leukemia. Blood. 2012;119(17):3917–3924.

16. Cooper TM, Sison EAR, Baker SD, et al. A phase 1 study of the CXCR4 antagonist plerixafor in combination with high-dose cytarabine and etoposide in children with relapsed or refractory acute leukemias or myelodysplastic syndrome: A Pediatric Oncology Experimental Therapeutics Investigators’ Consortium study (POE 10-03). Pediatr Blood Cancer. 2017;64(8).

17. Azab AK, Runnels JM, Pitsillides C, et al. CXCR4 inhibitor AMD3100 disrupts the interaction of multiple myeloma cells with the bone marrow microenvironment and enhances their sensitivity to therapy. Blood. 2009;113(18):4341–4351.

18. Chiang MY, Shestova O, Xu L, Aster JC, Pear WS. Divergent effects of supraphysiologic Notch signals on leukemia stem cells and hematopoietic stem cells. Blood. 2013;121(6):905–917.

19. Chiang MY, Xu ML, Histen G, et al. Identification of a conserved negative regulatory sequence that influences the leukemogenic activity of NOTCH1. Mol Cell Biol. 2006;26(16):6261–6271.

20. Khan AO, Rodriguez-Romera A, Reyat JS, et al. Human Bone Marrow Organoids for Disease Modeling, Discovery, and Validation of Therapeutic Targets in Hematologic Malignancies. Cancer Discov. 2023;13(2):364–385.

21. Olijnik AA, Rodriguez-Romera A, Wong ZC, et al. Generating human bone marrow organoids for disease modeling and drug discovery. Nat Protoc. 2024;19(7):2117–2146.

22. Dobin A, Davis CA, Schlesinger F, et al. STAR: ultrafast universal RNA-seq aligner. Bioinformatics. 2013;29(1):15–21.

23. Li B, Dewey CN. RSEM: accurate transcript quantification from RNA-Seq data with or without a reference genome. BMC Bioinformatics. 2011;12:323.

24. Benjamini Y, Hochberg Y. Controlling the False Discovery Rate: A Practical and Powerful Approach to Multiple Testing. Journal of the Royal Statistical Society: Series B (Methodological). 1995;57(1):289–300.

25. Kramer A, Green J, Pollard J, Jr., Tugendreich S. Causal analysis approaches in Ingenuity Pathway Analysis. Bioinformatics. 2014;30(4):523–530.

26. Bolte S, Cordelieres FP. A guided tour into subcellular colocalization analysis in light microscopy. J Microsc. 2006;224(Pt 3):213-232.

27. Liu Z, Chen S, Jin X, et al. Genome editing of the HIV co-receptors CCR5 and CXCR4 by CRISPR-Cas9 protects CD4(+) T cells from HIV-1 infection. Cell Biosci. 2017;7:47.

28. Sanjana NE, Shalem O, Zhang F. Improved vectors and genome-wide libraries for CRISPR screening. Nat Methods. 2014;11(8):783–784.

29. Shalem O, Sanjana NE, Hartenian E, et al. Genome-scale CRISPR-Cas9 knockout screening in human cells. Science. 2014;343(6166):84–87.

30. Reddy JK, Hashimoto T. Peroxisomal beta-oxidation and peroxisome proliferator-activated receptor alpha: an adaptive metabolic system. Annu Rev Nutr. 2001;21:193–230.

31. Winkler IG, Pettit AR, Raggatt LJ, et al. Hematopoietic stem cell mobilizing agents G-CSF, cyclophosphamide or AMD3100 have distinct mechanisms of action on bone marrow HSC niches and bone formation. Leukemia. 2012;26(7):1594–1601.

32. Yu M, Gang EJ, Parameswaran R, et al. AMD3100 sensitizes acute lymphoblastic leukemia cells to chemotherapy in vivo. Blood Cancer J. 2011;1(4):e14.

33. Welschinger R, Liedtke F, Basnett J, et al. Plerixafor (AMD3100) induces prolonged mobilization of acute lymphoblastic leukemia cells and increases the proportion of cycling cells in the blood in mice. Exp Hematol. 2013;41(3):293–302 e291.

34. Heredia JD, Park J, Brubaker RJ, Szymanski SK, Gill KS, Procko E. Mapping Interaction Sites on Human Chemokine Receptors by Deep Mutational Scanning. J Immunol. 2018;200(11):3825–3839.

35. Brelot A, Heveker N, Montes M, Alizon M. Identification of residues of CXCR4 critical for human immunodeficiency virus coreceptor and chemokine receptor activities. J Biol Chem. 2000;275(31):23736–23744.

36. Forster R, Kremmer E, Schubel A, et al. Intracellular and surface expression of the HIV-1 coreceptor CXCR4/fusin on various leukocyte subsets: rapid internalization and recycling upon activation. J Immunol. 1998;160(3):1522–1531.

37. Mills SC, Goh PH, Kudatsih J, et al. Cell migration towards CXCL12 in leukemic cells compared to breast cancer cells. Cell Signal. 2016;28(4):316–324.

38. Dunn KW, Kamocka MM, McDonald JH. A practical guide to evaluating colocalization in biological microscopy. Am J Physiol Cell Physiol. 2011;300(4):C723–742.

39. Olivas-Aguirre M, Pottosin I, Dobrovinskaya O. Mitochondria as emerging targets for therapies against T cell acute lymphoblastic leukemia. J Leukoc Biol. 2019;105(5):935–946.

40. Lee EA, Angka L, Rota SG, et al. Targeting Mitochondria with Avocatin B Induces Selective Leukemia Cell Death. Cancer Res. 2015;75(12):2478–2488.

41. Jitschin R, Hofmann AD, Bruns H, et al. Mitochondrial metabolism contributes to oxidative stress and reveals therapeutic targets in chronic lymphocytic leukemia. Blood. 2014;123(17):2663–2672.

42. Bodaar K, Yamagata N, Barthe A, et al. JAK3 mutations and mitochondrial apoptosis resistance in T-cell acute lymphoblastic leukemia. Leukemia. 2022;36(6):1499–1507.

43. Mondet J, Lo Presti C, Chevalier S, et al. Mitochondria in human acute myeloid leukemia cell lines have ultrastructural alterations linked to deregulation of their respiratory profiles. Exp Hematol. 2021;98:53–62 e53.

44. Xiao X, Yang J, Li R, et al. Deregulation of mitochondrial ATPsyn-beta in acute myeloid leukemia cells and with increased drug resistance. PLoS One. 2013;8(12):e83610.

45. Enzenmuller S, Niedermayer A, Seyfried F, et al. Venetoclax resistance in acute lymphoblastic leukemia is characterized by increased mitochondrial activity and can be overcome by co-targeting oxidative phosphorylation. Cell Death Dis. 2024;15(7):475.

46. Trombetti S, Cesaro E, Catapano R, et al. Oxidative Stress and ROS-Mediated Signaling in Leukemia: Novel Promising Perspectives to Eradicate Chemoresistant Cells in Myeloid Leukemia. Int J Mol Sci. 2021;22(5).

47. Chen C, Hao X, Lai X, et al. Oxidative phosphorylation enhances the leukemogenic capacity and resistance to chemotherapy of B cell acute lymphoblastic leukemia. Sci Adv. 2021;7(11).

48. Chen Y, Li J, Zhao Z. Redox Control in Acute Lymphoblastic Leukemia: From Physiology to Pathology and Therapeutic Opportunities. Cells. 2021;10(5).

49. Chen YF, Liu H, Luo XJ, et al. The roles of reactive oxygen species (ROS) and autophagy in the survival and death of leukemia cells. Crit Rev Oncol Hematol. 2017;112:21–30.

50. Geldenhuys WJ, Piktel D, Moore JC, et al. Loss of the redox mitochondrial protein mitoNEET leads to mitochondrial dysfunction in B-cell acute lymphoblastic leukemia. Free Radic Biol Med. 2021;175:226–235.

51. Skrtic M, Sriskanthadevan S, Jhas B, et al. Inhibition of mitochondrial translation as a therapeutic strategy for human acute myeloid leukemia. Cancer Cell. 2011;20(5):674–688.

52. Schimmer AD, Hedley DW, Penn LZ, Minden MD. Receptor-and mitochondrial-mediated apoptosis in acute leukemia: a translational view. Blood. 2001;98(13):3541–3553.

53. Zhou Y, Hileman EO, Plunkett W, Keating MJ, Huang P. Free radical stress in chronic lymphocytic leukemia cells and its role in cellular sensitivity to ROS-generating anticancer agents. Blood. 2003;101(10):4098–4104.

54. Pelicano H, Carney D, Huang P. ROS stress in cancer cells and therapeutic implications. Drug Resist Updat. 2004;7(2):97–110.

55. Basak NP, Banerjee S. Mitochondrial dependency in progression of acute myeloid leukemia. Mitochondrion. 2015;21:41–48.

56. Panina SB, Pei J, Kirienko NV. Mitochondrial metabolism as a target for acute myeloid leukemia treatment. Cancer Metab. 2021;9(1):17.

57. Jhas B, Sriskanthadevan S, Skrtic M, et al. Metabolic adaptation to chronic inhibition of mitochondrial protein synthesis in acute myeloid leukemia cells. PLoS One. 2013;8(3):e58367.

58. Fujimoto M, Hayashi T. New insights into the role of mitochondria-associated endoplasmic reticulum membrane. Int Rev Cell Mol Biol. 2011;292:73–117.

59. Jennings MJ, Hathazi D, Nguyen CDL, et al. Intracellular Lipid Accumulation and Mitochondrial Dysfunction Accompanies Endoplasmic Reticulum Stress Caused by Loss of the Co-chaperone DNAJC3. Front Cell Dev Biol. 2021;9:710247.

60. Wall CTJ, Lefebvre G, Metairon S, Descombes P, Wiederkehr A, Santo-Domingo J. Mitochondrial respiratory chain dysfunction alters ER sterol sensing and mevalonate pathway activity. J Biol Chem. 2022;298(3):101652.

61. van Vliet AR, Agostinis P. Mitochondria-Associated Membranes and ER Stress. Curr Top Microbiol Immunol. 2018;414:73–102.

62. Ribas V, Garcia-Ruiz C, Fernandez-Checa JC. Mitochondria, cholesterol and cancer cell metabolism. Clin Transl Med. 2016;5(1):22.

63. Brendolan A, Russo V. Targeting cholesterol homeostasis in hematopoietic malignancies. Blood. 2022;139(2):165–176.

64. Kalatskaya I, Berchiche YA, Gravel S, Limberg BJ, Rosenbaum JS, Heveker N. AMD3100 is a CXCR7 ligand with allosteric agonist properties. Mol Pharmacol. 2009;75(5):1240–1247.

