## Supplemental Materials for "PAMD-Ch17, a Polymeric Analog of Plerixafor, Induces Mitochondrial Dysfunction in T-ALL Cells Independent of CXCR4"

### SUPPLEMENTAL FIGURE LEGENDS

**Supplemental Figure 1. AMD3100 does not have anti-leukemic effects.** Graph of relative metabolic activity for the indicated human AML and ALL cell lines treated with 5μM AMD3100 for 72 hours. N=3.

**Supplemental Figure 2. PAMD-Ch17 blocks the CXCR4 receptor.** Representative flow cytometry plots of anti-CXCR4 12G5 and 2B11 staining in Jurkat cells treated with either 0.6μM AMD3100, PAMD-Ch17, or untreated for 24 hours.

**Supplemental Figure 3. Validation of *CXCR4* KO Jurkat cells.** A) Representative flow cytometry plots of anti-CXCR4 12G5 and 2B11 staining in Jurkat cells of the indicated genotypes. B) Immunofluorescence microcopy images of Jurkat cells of the indicated genotypes stained with anti-CXCR4 UMB2 and 2B11. C) Graph of relative migration in response to CXCL12 in Jurkat cells of the indicated genotypes. N=3. Scale bar is 10um. * $p \leq 0.05$.

**Supplemental Figure 4. PAMD-Ch17 dysregulates genes different from that of *CXCR4* KO.** A) Venn diagram of the differentially expressed genes in *CXCR4* KO-2 and PAMD-Ch17 treated, as compared to wild type Jurkat cells. B) Ingenuity Pathway analysis of the pathways associated with the differentially expressed genes in *CXCR4* KO-2 cells. N=3.

**Supplemental Figure 5. PAMD-Ch17 colocalizes with the mitochondria.** Representative confocal images of Jurkat cells treated with fluorescently tagged PAMD-Ch17 (fPAMD) at 0.6uM (A) or 1.2uM (B) for the indicated time before staining with Mitotracker Deep Red and DAPI. N=3. Scale bar is 10um.

**Supplemental Figure 6. PAMD-Ch17's effects on mouse primary T-ALL cells.** A) Representative flow cytometry plots and graph of the percentage of primary Notch1 ΔEGFΔLNRΔP expressing T-ALL cells from 4 independent mice that are $CD8^+$. B) Representative flow cytometry plots and graph of the percentage (%) of Mitosox+ and C) 7-$AAD^+$ mouse bone marrow cells treated with 1.2uM PAMD-Ch17 for the indicated time. UT= untreated, T= PAMD-Ch17 treated. N=3 healthy mouse bone marrow samples.

**Supplemental Figure 7. Human primary T-ALL cells engraftment in human bone marrow organoids.** Graph of the percentage of human bone marrow organoids cells that are $CellTrace^+$ leukemia cells for organoids engrafted with the indicated patient T-ALL sample . Untreated organoids of the 4 and 24 hour conditions shown. 5-7 engrafted organoids were combined for each replicate. Each condition has 3 replicates.

SFig.1

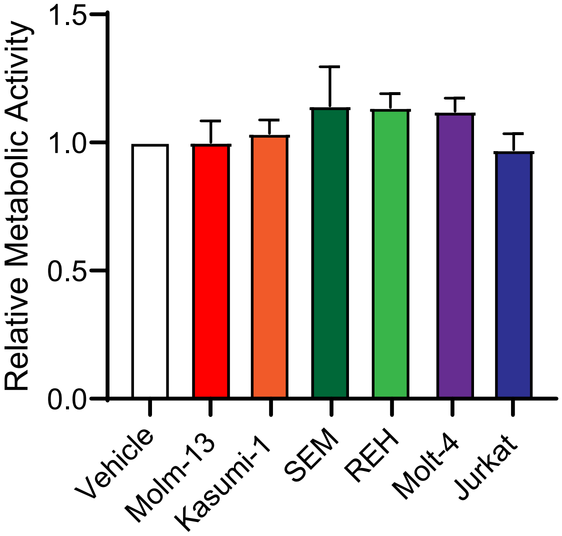

Relative Metabolic Activity
1.5
1.0
0.5
0.0
Vehicle
Molm-13
Kasumi-1
SEM
REH
Molt-4
Jurkat

SFig.2

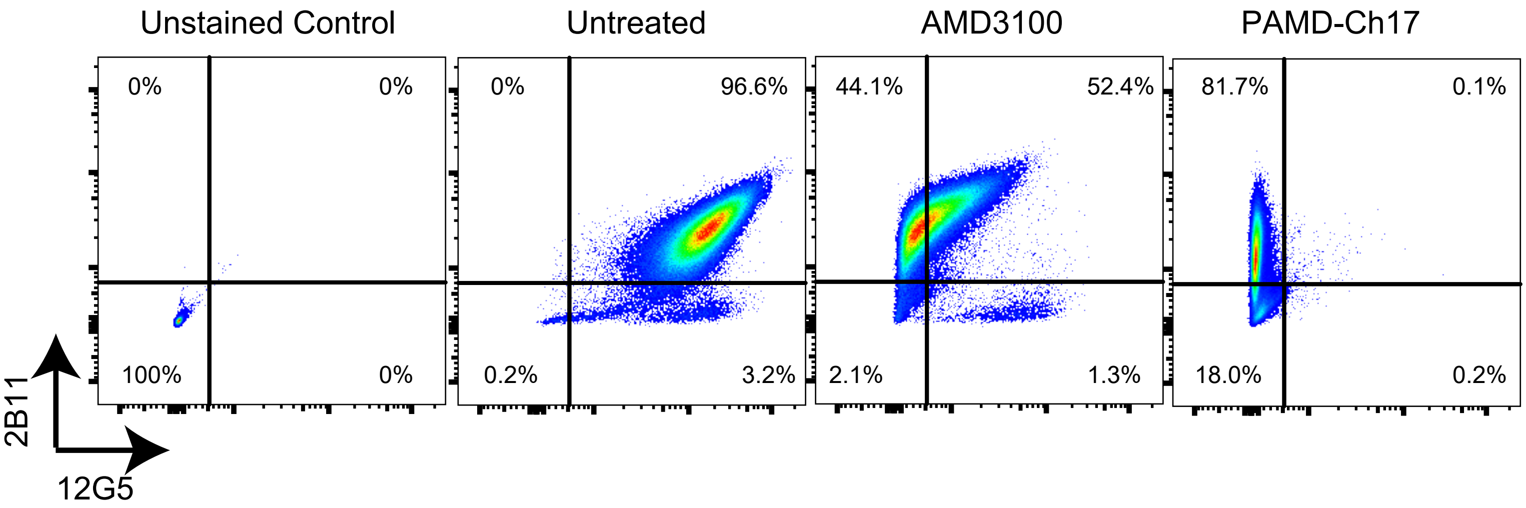
Unstained Control
Untreated
AMD3100
PAMD-Ch17
0%
0%
100%
0%
0%
96.6%
0.2%
3.2%
44.1%
52.4%
2.1%
1.3%
81.7%
0.1%
18.0%
0.2%
2B11
12G5

SFig.3

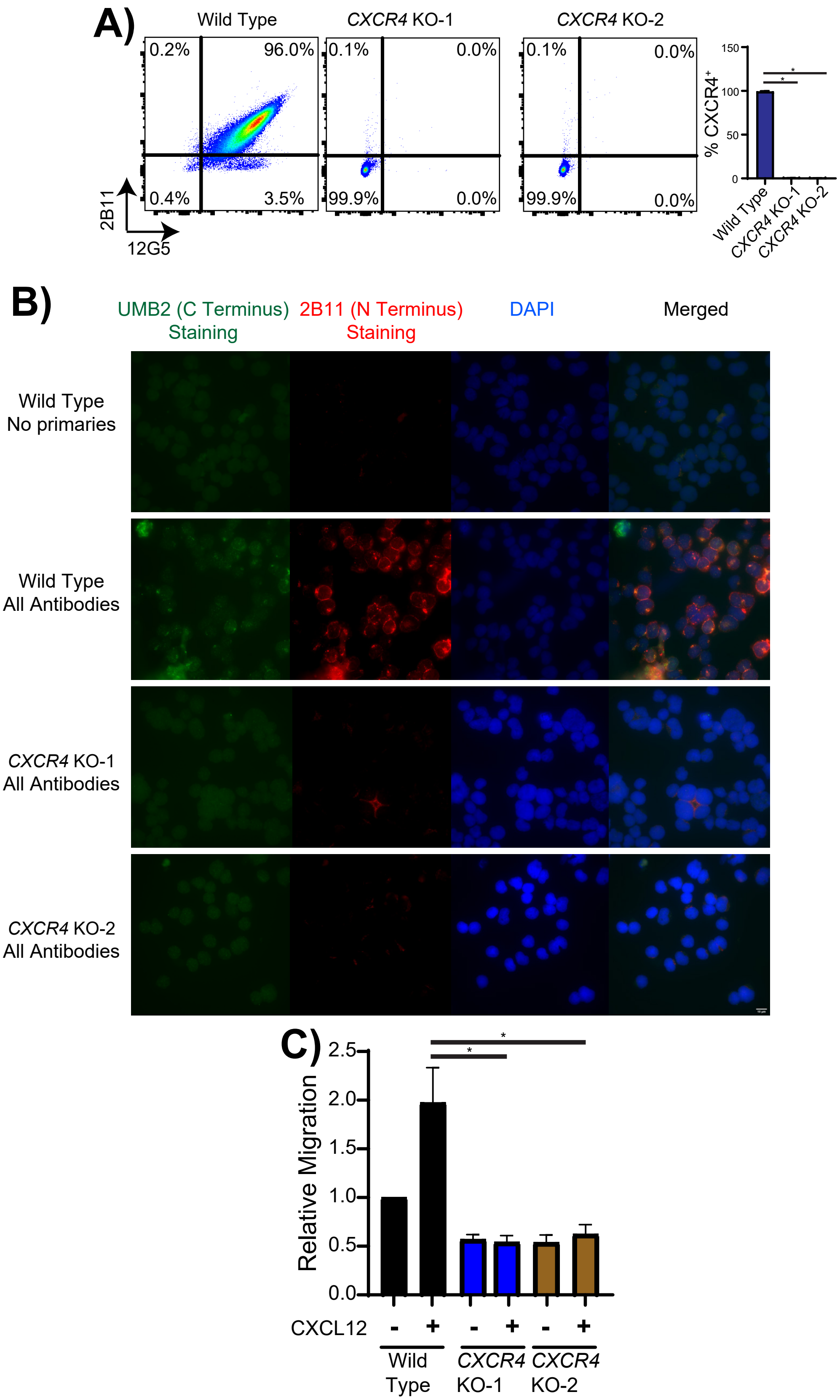
A)
Wild Type
CXCR4 KO-1
CXCR4 KO-2
0.2%
96.0%
0.4%
3.5%
0.1%
0.0%
99.9%
0.0%
0.1%
0.0%
99.9%
0.0%
2B11
12G5
% CXCR4+
150
100
50
0
*
*
Wild Type
CXCR4 KO-1
CXCR4 KO-2
B)
UMB2 (C Terminus) Staining
2B11 (N Terminus) Staining
DAPI
Merged
Wild Type No primaries
Wild Type All Antibodies
CXCR4 KO-1 All Antibodies
CXCR4 KO-2 All Antibodies
C)
Relative Migration
2.5
2.0
1.5
1.0
0.5
0.0
*
*
CXCL12
-
+
-
+
-
+
Wild Type
CXCR4 KO-1
CXCR4 KO-2

SFig.4

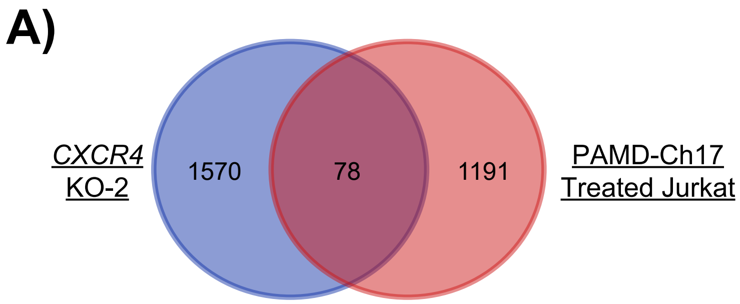
A)
CXCR4
KO-2
1570
78
1191
PAMD-Ch17
Treated Jurkat

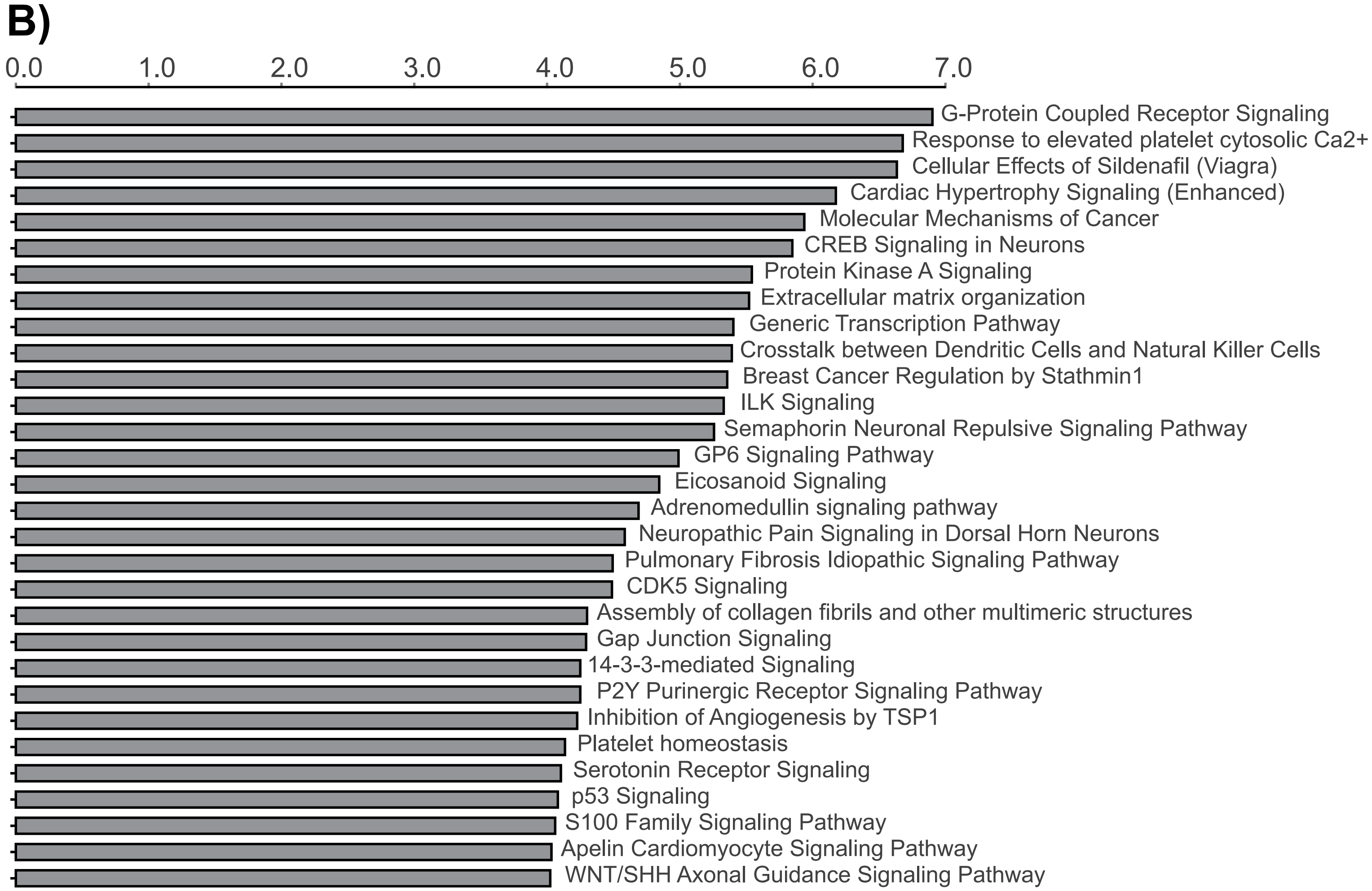
B)
0.0
1.0
2.0
3.0
4.0
5.0
6.0
7.0
G-Protein Coupled Receptor Signaling
Response to elevated platelet cytosolic Ca2+
Cellular Effects of Sildenafil (Viagra)
Cardiac Hypertrophy Signaling (Enhanced)
Molecular Mechanisms of Cancer
CREB Signaling in Neurons
Protein Kinase A Signaling
Extracellular matrix organization
Generic Transcription Pathway
Crosstalk between Dendritic Cells and Natural Killer Cells
Breast Cancer Regulation by Stathmin1
ILK Signaling
Semaphorin Neuronal Repulsive Signaling Pathway
GP6 Signaling Pathway
Eicosanoid Signaling
Adrenomedullin signaling pathway
Neuropathic Pain Signaling in Dorsal Horn Neurons
Pulmonary Fibrosis Idiopathic Signaling Pathway
CDK5 Signaling
Assembly of collagen fibrils and other multimeric structures
Gap Junction Signaling
14-3-3-mediated Signaling
P2Y Purinergic Receptor Signaling Pathway
Inhibition of Angiogenesis by TSP1
Platelet homeostasis
Serotonin Receptor Signaling
p53 Signaling
S100 Family Signaling Pathway
Apelin Cardiomyocyte Signaling Pathway
WNT/SHH Axonal Guidance Signaling Pathway

SFig.5

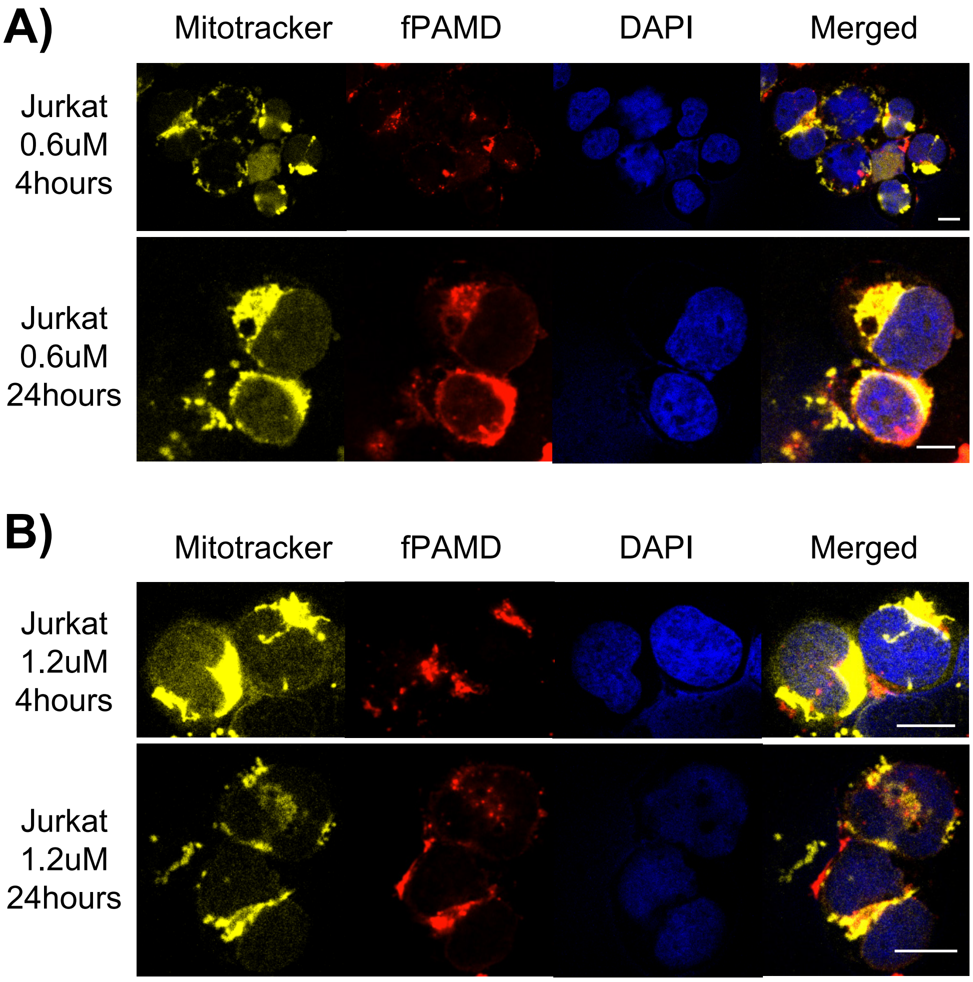
A)
Mitotracker
fPAMD
DAPI
Merged
Jurkat
0.6uM
4hours
Jurkat
0.6uM
24hours
B)
Mitotracker
fPAMD
DAPI
Merged
Jurkat
1.2uM
4hours
Jurkat
1.2uM
24hours

SFig.6

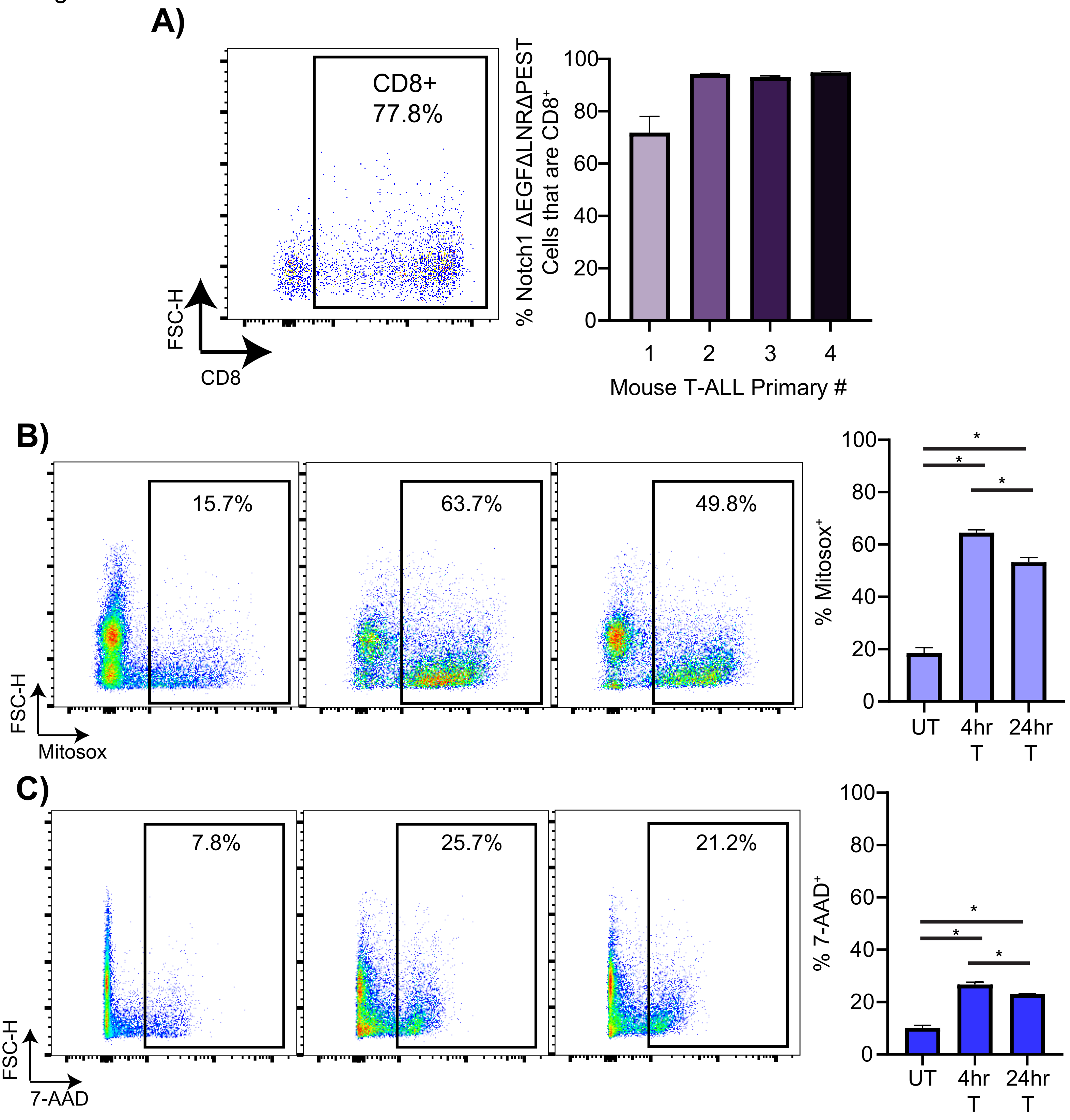

A)
CD8+
77.8%
FSC-H
CD8
% Notch1 ΔEGFΔLNRΔPEST Cells that are CD8+
100
80
60
40
20
0
1
2
3
4
Mouse T-ALL Primary #
B)
15.7%
63.7%
49.8%
FSC-H
Mitosox
% Mitosox+
*
UT
4hr T
24hr T
C)
7.8%
25.7%
21.2%
FSC-H
7-AAD
% 7-AAD+

SFig.7

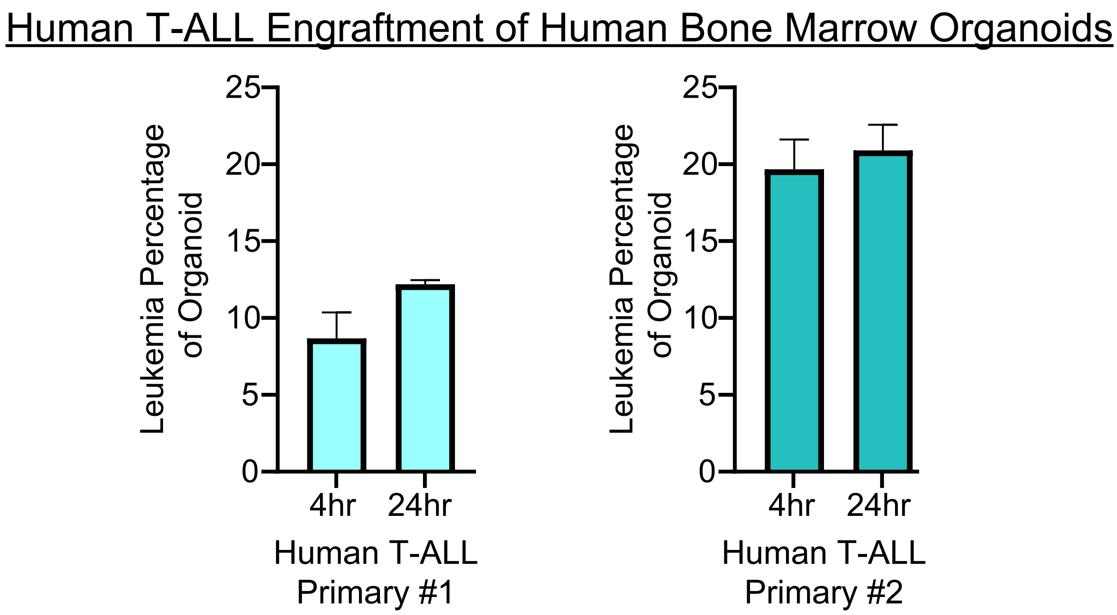
Human T-ALL Engraftment of Human Bone Marrow Organoids
Leukemia Percentage of Organoid
25
20
15
10
5
0
4hr
24hr
Human T-ALL Primary #1
Leukemia Percentage of Organoid
25
20
15
10
5
0
4hr
24hr
Human T-ALL Primary #2
